# MTERF1 loss buffers against pathogenic mtDNA deletions through transcriptional regulation

**DOI:** 10.64898/2026.07.16.738964

**Authors:** Tamar Kavlashvili, Elim Choi, Ruobing Cui, Juan C. Landoni, Simon Zhang, Stefan Isaac, Stirling Churchman, Suliana Manley, Craig Thompson, Agnel Sfeir

## Abstract

Large-scale mitochondrial DNA (mtDNA) deletions cripple oxidative phosphorylation once they exceed a critical heteroplasmy threshold, causing incurable mitochondrial pathologies. Using a genome-wide CRISPR/Cas9 screen in an engineered human cell line carrying a large-scale mtDNA deletion at high heteroplasmy, we identified mitochondrial transcription termination factor 1 (MTERF1) as a suppressor of the heteroplasmy burden. Loss of MTERF1 restored mitochondrial function and increased cellular proliferation in cells with a mtDNA deletion burden exceeding the pathogenic threshold, without altering heteroplasmy or mtDNA copy number. MTERF1 binds wild-type and deletion-bearing mitochondrial genomes indiscriminately at a site downstream of the ribosomal RNA genes and curbs transcription. Relieving this constraint broadly increased OXPHOS transcripts, thereby eliciting more respiratory output from the residual wild-type genomes. Notably, the buffering effect of MTERF1 loss extended beyond mtDNA deletions. In a counter-screen, MTERF1 loss could also restore respiratory growth in cells depleted of nuclear-encoded mitochondrial genes such as *OPA1* and *COX5A*. Together, these findings indicate that by relieving a transcriptional constraint, MTERF1 loss compensates for reduced genome dosage, defining a strategy to enhance residual mitochondrial function in mtDNA deletion disorders and related conditions.

## Introduction

Mitochondria carry their own compact, multi-copy genome encoding 13 essential subunits of the oxidative phosphorylation (OXPHOS) machinery, along with the ribosomal and transfer RNAs required for their synthesis. These mitochondria-encoded subunits combine with a larger complement of nuclear-encoded proteins to assemble the respiratory chain complexes that drive cellular ATP synthesi^1^. Pathogenic mtDNA alterations, including point mutations and large-scale deletions, are a leading cause of inherited metabolic disease, affecting roughly 1 in 4,300 individuals^2^. Because each cell carries hundreds to thousands of mtDNA copies, mutant and wild-type genomes can coexist in a state of heteroplasmy, in which wild-type copies buffer the defect until the mutant fraction exceeds ∼75%, beyond which cells lose respiratory capacity^3,4^. The most intractable of these lesions are large-scale deletions, including the mitochondrial common deletion, a prevalent mutation that removes a 4977 bp region coding for multiple OXPHOS genes and tRNAs. At high heteroplasmy, such deletions drive devastating disorders, including Pearson and Kearns-Sayre syndromes and progressive external ophthalmoplegia (PEO)^5-7^. mtDNA deletions can arise sporadically or be linked to mutations in nuclear-encoded components of mtDNA replication machinery, including DNA polymerase gamma (POLG) and the helicase Twinkle^4,5,8^. Once formed, deletion-bearing genomes undergo clonal expansion in post-mitotic tissues such as skeletal muscle and brain, where they accumulate progressively with age and contribute to sarcopenia and selective neuronal vulnerability^9-11^. Yet patients with mitochondrial deletion syndromes remain without effective treatment. Nuclease- and base-editing strategies that shift heteroplasmy are being tested for mtDNA point mutations, but don’t apply to deletions^12-14^. Other emerging therapies in clinical testing, including POLG reactivation by molecules such as PZL-A and nucleotide supplementation to replenish dNTP pools, target deficiencies associated with impaired mtDNA maintenance but cannot reverse deletions already fixed at pathogenic loads.

Uncovering nuclear genes that mitigate the cellular consequences of mtDNA deletions, whether by driving selection against deletion-bearing genomes or by buffering bioenergetic defect, has been hampered by the lack of isogenic human cell models in which deletion load can be precisely controlled. To address this, we recently established such a model by reconstituting end-joining in human mitochondria (Mito-EJ) in cells expressing a mitochondrially targeted restriction enzyme (mito-ScaI), generating defined double-strand breaks across a ∼3.5 kb fragment that largely overlaps with the common deletion. This approach produced an isogenic ARPE-19 panel carrying mtDNA deletions (mtDNA^ΔScaI^) across a full spectrum of heteroplasmy^15^. Using this model, we found that cells exceeding ∼75% heteroplasmy undergo progressive mitochondrial deterioration, with suppressed oxygen consumption, loss of OXPHOS complex subunits, altered mitochondrial metabolite pools, and a broad transcriptional response reflecting metabolic stress. Forced to rely on respiration when cultured in galactose, cells with >75% heteroplasmy fail to survive, thereby establishing a defined platform to interrogate which nuclear genes modulate cellular fitness under high deletion burden.

Here, we exploit this isogenic mtDNA^ΔScaI^ cell panel to conduct a genome-wide CRISPR screen for nuclear genes whose loss suppresses growth defects and respiratory failure in high-deletion cells. The top hit, MTERF1, a mitochondrial transcription termination factor of previously unknown relevance to disease, emerged as a potent suppressor of deletion-based defects. Its loss restored respiration and growth in cells above the pathogenic threshold without altering heteroplasmy or mtDNA copy number. Rather than altering genome composition, MTERF1 loss increases mitochondrial transcriptional output, thereby extracting more OXPHOS capacity from the residual wild-type genomes. In high heteroplasmy cells. A genome-wide counter-screen showed that this benefit generalizes across diverse metabolic defects, including loss of OPA1, mutated in autosomal dominant optic atrophy^16,17^, and COX5A, loss of which causes complex IV deficiency^18,19^. MTERF1, therefore, acts as a brake on transcriptional reserve, and its release offers a mutation-agnostic route to boost residual mitochondrial function.

## Results

### A genome-wide CRISPR/Cas9 screen identifies MTERF1 as a suppressor of respiratory failure at high mtDNA deletion heteroplasmy

Using the two-component Mito-EJ system and mitochondrial-targeted ScaI^15^, we generated clonally derived ARPE-19 cells carrying mtDNA^ΔScaI^ deletion at 95% and 100% heteroplasmy (**Fig S1A-C**). In glucose medium, which permits glycolytic ATP production, high-heteroplasmy cells proliferate comparably to wild-type cells (**Fig. 1A, top**). In galactose medium, which enforces OXPHOS dependency, both high heteroplasmy cell lines fail to proliferate (**Fig. 1A, bottom**). We exploited this conditional lethality to perform a genome-wide CRISPR/Cas9 screen for nuclear genes whose loss confers a selective advantage under OXPHOS dependency.

**Figure 1:**
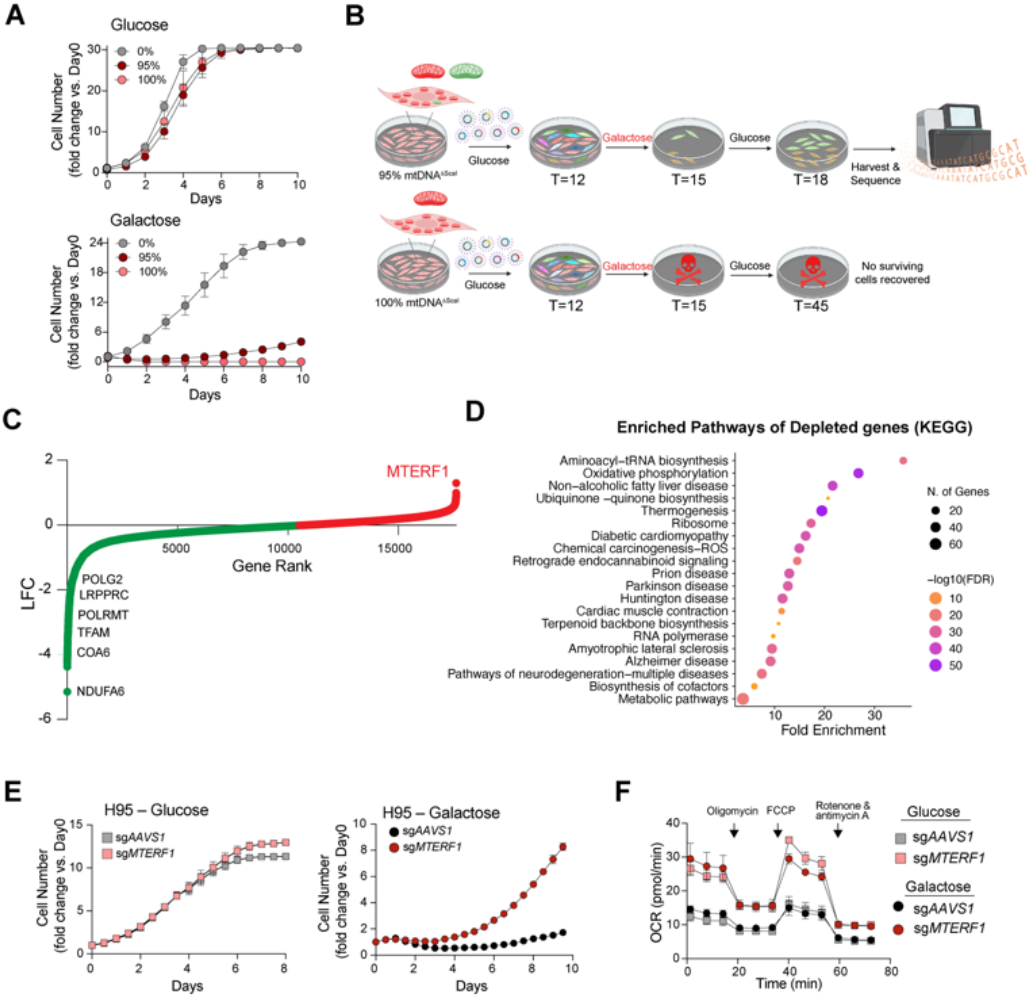
Genome-wide CRISPR Screen Identifies MTERF1 as a suppressor of mtDNA deletion at high heteroplasmy. **(A)** Growth curves of cells harboring the indicated mtDNA^ΔScaI^ heteroplasmy (0%, 95%, 100%) in 25 mM glucose (top) or 10 mM galactose (bottom) medium. Data presented as fold change in cell number relative to Day 0. **(B)** Schematic of genome-wide CRISPR screen experimental outline. Cells harboring 95% mtDNA^ΔScaI^ heteroplasmy or 100% mtDNA^ΔScaI^ homoplasmy cells were transduced with TKOv3 sgRNA library, selected with 5 µg/mL puromycin for 48 h beginning 24 h post-infection, and expanded in glucose medium for 12 days. At T = 12, cells were switched to galactose medium for 3 days (T = 15). Surviving cells from the 95% heteroplasmy arm were recovered by an additional 3 days of glucose culture (T = 18), harvested, and subjected to next-generation sequencing (NGS) to determine sgRNA abundance. No surviving cells were recovered from the 100% homoplasmy arm. **(C)** CRISPR screen data were analyzed using the MAGeCK algorithm. Each point represents a gene, ranked by log2 fold change (LFC). Select depleted genes (essential under OXPHOS stress) and the top enriched gene, *MTERF1*, are labeled. **(D)** Dot plot of selected KEGG pathways enriched among negatively selected genes (FDR < 0.05). Dot size indicates the number of genes per pathway; color indicates −log10(FDR). No biological pathways were significantly enriched among positively selected genes at an FDR threshold of 0.05. **(E)** Growth curves of sg*AAVS1*- or sg*MTERF1*-targeted 95% mtDNA^ΔScaI^ cells cultured in glucose (left) or galactose (right) medium. Data are presented as fold change in cell number relative to day 0. **(F)** Oxygen consumption rate (OCR) measured by Seahorse extracellular flux analysis in sg*AAVS1*- or sg*MTERF1*-targeted cells cultured in glucose (squares) or galactose (circles) medium. Arrows indicate sequential injections of oligomycin, FCCP, and rotenone plus antimycin A. Cell lines analyzed: 95% mtDNA^ΔScaI^ sgAAVS1 and sgMTERF1.

Cells carrying 95% or 100% mutant load were transduced with the genome-wide TKOv3 library, selected with puromycin, expanded in glucose for 12 days, and then shifted to galactose for three days to enforce OXPHOS reliance. Surviving cells were then returned to glucose for three days to recover before the population was harvested and subjected to next-generation sequencing for sgRNA enrichment and depletion analysis using MAGeCK ^20^ (**Fig. 1B**). The CRISPR/Cas9 screen achieved strong library coverage, sequencing depth, and essential-gene dropout using BAGEL analysis (**Fig. S1D– G and Table S1**), confirming robust performance. Cells that are homoplasmic for mutant mtDNA (100% mtDNA^ΔScaI^) failed to survive the metabolic challenge regardless of perturbation, indicating that wild-type genomes are strictly required for fitness buffering (**Fig. 1B**). By contrast, a small fraction of cells with 95% heteroplasmy survived and were then carried forward for analysis. Analysis of sgRNA representation showed depletion of well-established mitochondrial maintenance factors, including POLG, LRPPRC, POLRMT, and TFAM, confirming that the screen recovers genes essential for OXPHOS function (**Fig. 1C**). Furthermore, pathway analysis of depleted genes revealed strong enrichment for aminoacyl-tRNA biosynthesis and oxidative phosphorylation, confirming that the screen is selectively reporting on mitochondrial metabolic function (**Fig. 1D, Table S2**). Whereas depletion of essential mitochondrial genes was broad, enrichment was remarkably sparse, dominated by a single outlier, MTERF1 (mitochondrial transcription termination factor 1), loss of which conferred the strongest selective advantage under respiratory stress at high deletion burden (**Fig. 1C, Table S1**).

To validate that MTERF1 loss underlies the recovery of respiratory growth in high-heteroplasmy cells, we depleted the gene with independent sgRNAs in mtDNA^ΔScaI^ lines with 95% heteroplasmy (**Fig. S1H**). Cells expressing a control sgRNA targeting the safe-harbor *AAVS1* locus proliferated normally in glucose but uniformly failed in galactose, faithfully recapitulating the deletion-driven respiratory defect. MTERF1 knockout reversed this phenotype, restoring proliferation in galactose without affecting growth in glucose (**Fig. 1E**). This proliferative rescue tracked with improved respiration. Specifically, Seahorse analysis showed that MTERF1 loss raised oxygen consumption rates in 95% heteroplasmy cells under both glucose and galactose conditions (**Fig. 1F**), establishing that the benefit operates at the level of mitochondrial bioenergetics.

### MTERF1 loss restores mitochondrial architecture and respiratory capacity without altering heteroplasmy or mtDNA copy number

MTERF1 belongs to the vertebrate MTERF family^21^ with a known function as a mitochondrial transcription termination factor. It binds with high sequence specificity to a 28-bp site within the tRNALeu (UUR) gene, immediately downstream of the rRNA genes, where it bends, unwinds, and base-flips its DNA target through three stacking residues (R162, F243, Y288)^22,23^. MTERF1 was originally proposed to terminate heavy-strand transcription, delimiting a transcriptional unit that favors high-level synthesis of ribosomal RNAs (rRNA)^24^. This model was subsequently challenged *in vitro*, where MTERF1 was found to terminate transcription with strong polarity, blocking LSP-but not HSP-initiated transcripts, thereby preventing interference with the rDNA locus as well as the control region ^25,26^. *In vivo*, however, *Mterf1* knockout mice are viable and phenotypically normal, with unchanged rRNA levels but slightly reduced 7S RNA levels, suggesting that MTERF1 is dispensable for rRNA expression under physiological conditions^27^. It was unknown whether any of these activities could buffer the pathogenicity of mtDNA deletions.

To define the heteroplasmy range over which MTERF1 loss is beneficial and exclude a clonal artifact, we extended our analysis to additional mtDNA^ΔScaI^ lines spanning a full range of deletion loads. MTERF1 depletion in wild-type ARPE-19 cells had no detectable effect on respiration (Fig. 2A), establishing that MTERF1 loss is well tolerated in a healthy mitochondrial background.

**Figure 2:**
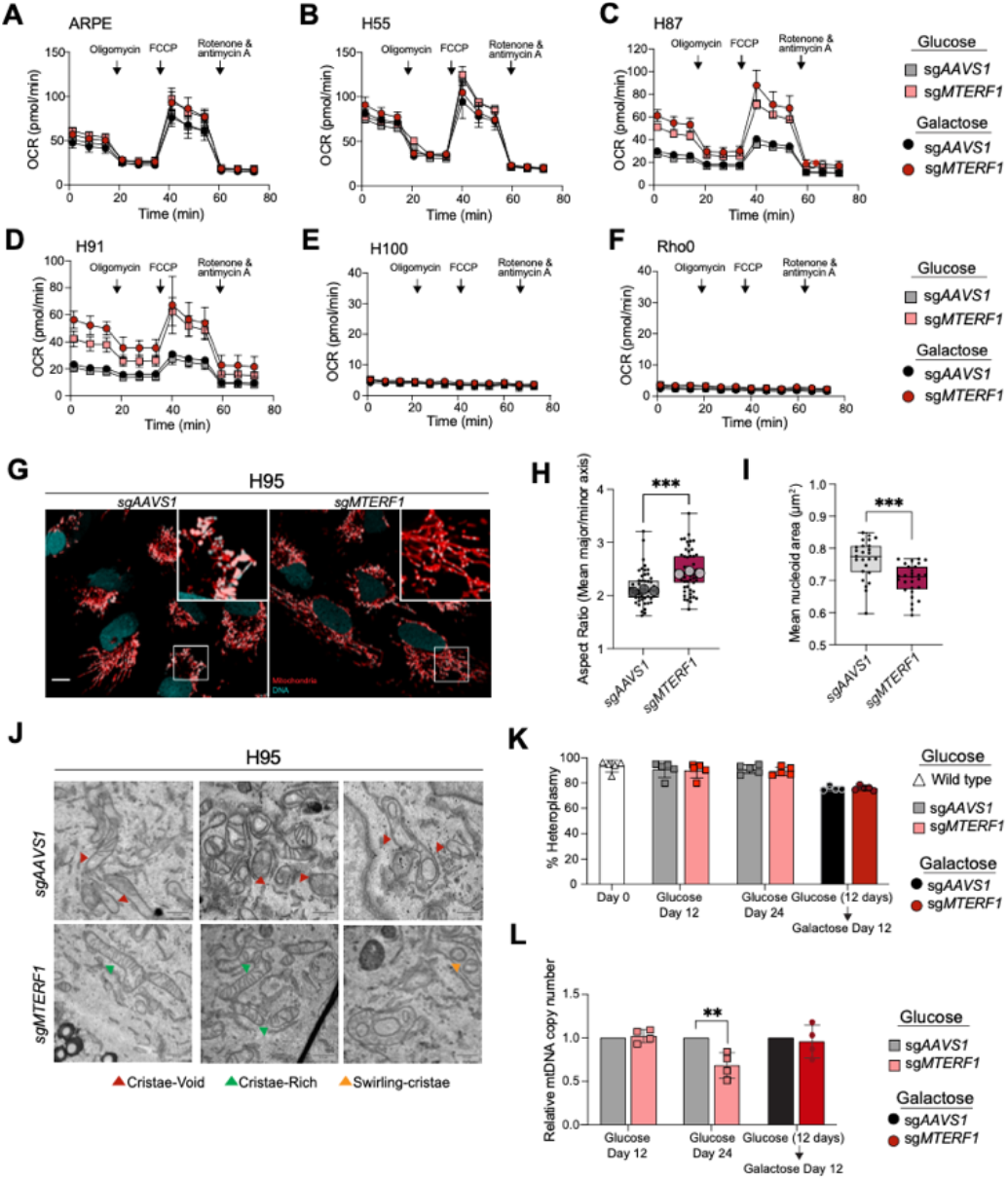
MTERF1 loss suppressed growth defect under OXPHOS stress without changing heteroplasmy. **(A**) Oxygen consumption rate (OCR) measured by Seahorse extracellular flux analysis in sgAAVS1- or sgMTERF1-targeted cells cultured in glucose (squares) or galactose (circles) medium. Arrows indicate sequential injections of oligomycin, FCCP, and rotenone plus antimycin A. Cell lines analyzed: (A) wild-type ARPE-19. (B) 55% mtDNA^ΔScaI^,(C) 87% mtDNA^ΔScaI^, (D) 91% mtDNA^ΔScaI^, (E) 100% mtDNA^ΔScaI^ and (F) ρ0 ARPE-19. **(G)** Representative live imaging of mitochondria (TMRE, red) and DNA (SYBR gold, teal), the latter staining both the nucleus and mtDNA nucleoids within mitochondria in sgAAVS1 and sgMTERF1 95% mtDNA^ΔScaI^ cell lines. Teal arrowhead highlights clustered. Scale bar, 10 µm. **(H)** Superplot of mean aspect ratios (major/minor axis) of segmented mitochondria per-cell. Small dots represent values per individual cell, large outlined circles represent the mean of each biological replicate (N = 3 independent experiments). Boxplots show median and interquartile range, p value from an unpaired Wilcox test (^***^p < 0.001). **(I)** Mean area of segmented nucleoids per cell. Boxplots show median and interquartile range, p value from an unpaired Wilcox test (^***^p < 0.001). **(J)** Electron micrographs of sg AAVS1 and sgMTERF1 95% mtDNA^ΔScaI^ cell lines. **(K)** Quantitative PCR (qPCR) analysis of mtDNA^ΔScaI^ levels sgAAVS1- or sgMTERF1-targeted 95% mtDNA^ΔScaI^ cells at day 0, after 12 or 24 days of glucose culture, or after 12 days of glucose followed by 12 days of galactose culture (24 days total). **(L)** qPCR quantification of relative mtDNA copy number in sgAAVS1- or sgMTERF1-targeted 95% mtDNA^ΔScaI^ cell lines at the same time points as in (I).

Similarly, no proliferative or respiratory benefit was observed in cells at 55% heteroplasmy under either glucose or galactose conditions (**Fig. 2B, S2A**), indicating that the effect is specific to high deletion burden. By contrast, MTERF1 loss robustly rescued proliferation and respiration in independent lines at 87% and 91% heteroplasmy (**Fig. 2C-D and Fig. S2B-C**), confirming that the benefit is reproducible across clones and emerges specifically above the pathogenic threshold. Crucially, neither growth nor respiration was rescued in homoplasmic (100%) deletion cells or in ρ^0^ cells devoid of mtDNA (**Fig. 2E-F**), establishing that the rescue requires residual wild-type template. This dependence was also reflected in glycolytic output. Lactate secretion, a readout of OXPHOS insufficiency, rose with increasing heteroplasmy, but MTERF1 loss lowered lactate in H95 cells below the threshold levels while H100 cells remained elevated (**Fig. S2D**). Together, these results establish that MTERF1 loss confers respiratory and proliferative benefits contingent on both a high deletion burden and the presence of wild-type genomes.

The functional rescue was mirrored by a striking reorganization of mitochondrial architecture. In control sgAAVS1 cells, mitochondria appeared fragmented, and mtDNA was irregularly distributed, with poorly organized nucleoid morphology. Loss of MTERF1 transformed this landscape: live imaging of 95% heteroplasmy cells revealed a shift toward a more interconnected, elongated mitochondrial network with nucleoids significantly more dispersed and uniform in size, as quantified by increased mitochondrial aspect ratio and reduced nucleoid size (**Fig. 2G–I**). Ultrastructural analysis by electron microscopy revealed the basis of this improvement at the organelle level. Control cells with 95% heteroplasmy were characterized by swollen, cristae-void, and cristae-poor mitochondria, hallmarks of bioenergetic failure. In contrast, MTERF1-depleted cells harbored a markedly higher occurrence of lamellar cristae ultrastructure, typical of healthy organelles (**Fig. 2J and Fig. S2E**). Given that high heteroplasmy triggers apoptosis in these cells^15^, the restored cristae architecture upon MTERF1 loss may reinforce cytochrome c sequestration, thereby prevent apoptosis and allowing continued proliferation^28^.

We next sought to understand the mechanism by which MTERF1 loss rescues high-heteroplasmy cells. We have previously shown that prolonged galactose culture drives a gradual reduction in heteroplasmy, consistent with selection against deletion-bearing genomes under respiratory stress^15^. We therefore hypothesized that MTERF1 loss might accelerate this process, shifting genome composition in favor of wild-type mtDNA. However, qPCR quantification of mutant-to-wild-type genome ratios revealed no difference between sgMTERF1 and control cells at 95% heteroplasmy across all glucose and galactose time points. These results suggest that the deletion fraction was neither cleared nor diluted under either condition (**Fig. 2K**). The same was true at 55% heteroplasmy (**Fig. S2F**), confirming that MTERF1 does not alter genome composition regardless of deletion load. Total mtDNA copy number was likewise unchanged, indicating that the restored respiratory capacity does not arise from increased genome abundance (**Fig. 2L and S2G)**. Together, these data indicate that MTERF1 loss rescues mitochondrial function and ultrastructure in high-heteroplasmy cells without altering mitochondrial genome composition or abundance.

### MTERF1 loss derepresses mitochondrial transcription to compensate for mtDNA deletions

To understand how MTERF1 loss compensates for the metabolic defect imposed by the ScaI deletion, we profiled gene expression by RNA-seq. Analysis of the nuclear transcriptome revealed minimal changes in nuclear gene expression (**Fig. S3A-B**) between MTERF1-knockout and control cells, regardless of heteroplasmy load, pointing to a mitochondria-autonomous mechanism for buffering against mutant genomes. By contrast, loss of MTERF1 led to significant upregulation of mitochondrial transcripts in both the 55% and 95% mtDNA^ΔScaI^ backgrounds. The notable exceptions were the 12S and 16S ribosomal RNAs, each being reduced by approximately twofold, consistent with the established role of MTERF1 in rRNA termination (**Fig. 3A and Fig. S3A–D, Tables S3–4**)^24^. A similar transcript profile was seen in 100% mtDNA^ΔScaI^ cells, indicating that outside the deleted region the genome is transcribed comparably and that these transcripts derive from both wild-type and deleted genomes (**Fig. 3A**). The 7S RNA, a non-coding transcript that inhibits mitochondrial transcription by inducing POLRMT dimerization, thereby sequestering the polymerase POLRMT in a catalytically inactive state^26^, was reduced in MTERF1-knockout cells (Fig. 3B, Fig. S3E). This reduction could potentially account for the broad increase in mitochondrial transcript levels observed upon MTERF1 depletion. We next asked whether the increase in transcript abundance was reflected at the protein level. Western blotting across a glucose-to-galactose time course showed that both mtDNA-encoded (COXII) and nuclear-encoded (NDUFB8, SDHB, UQCRC2, ATP5A) respiratory chain subunits accumulated to higher levels upon MTERF1 loss, in high heteroplasmy cells (**Fig. 3C-D and Fig. S3F**).

**Figure 3:**
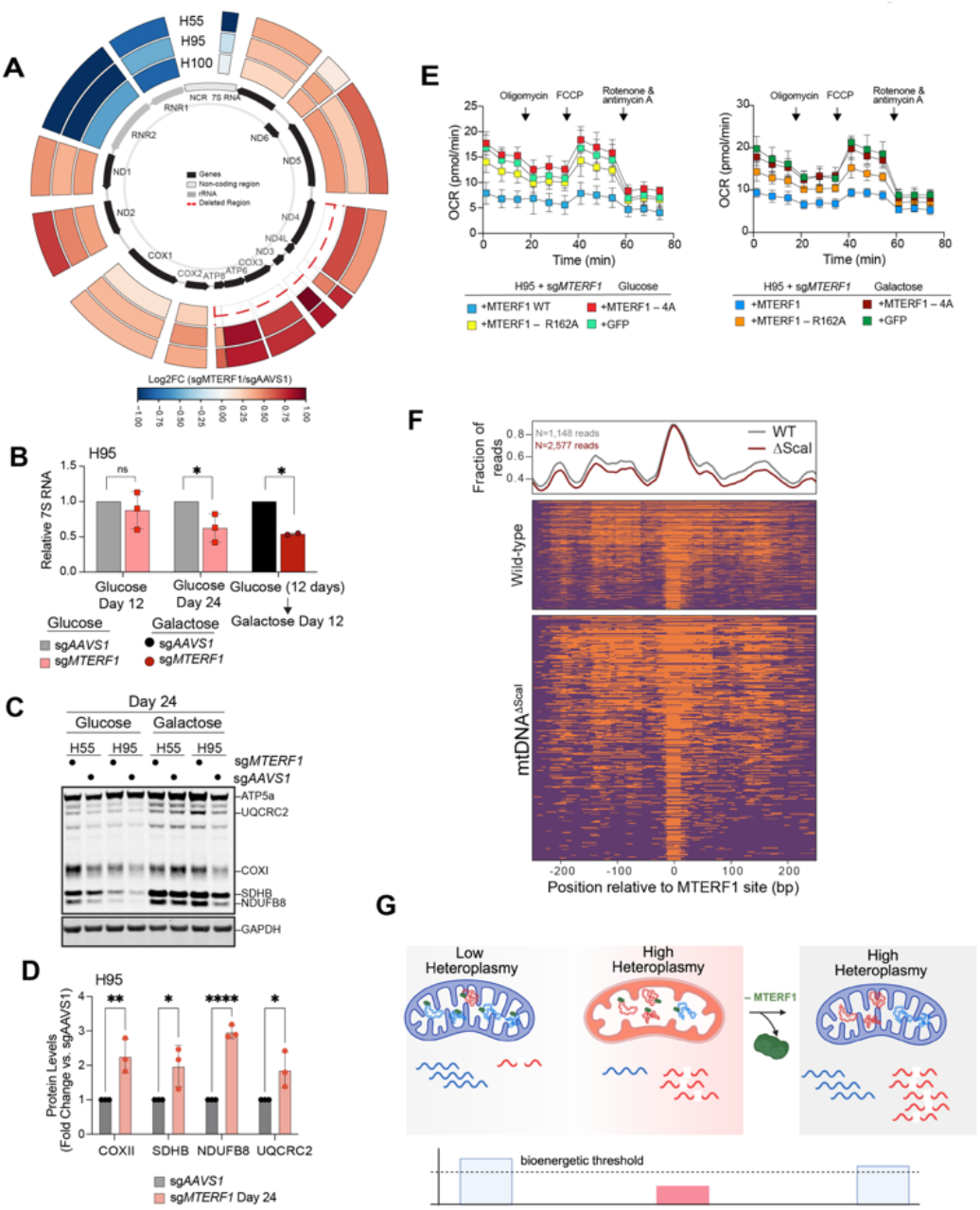
MTERF1 loss alleviates high heteroplasmy defect by increasing mtRNA transcript abundance. **(A)** RNA-sequencing analysis of mitochondrial transcript levels in sg*AAVS1*- or sg*MTERF1*-targeted 100%, 95% and 55% mtDNA^ΔScaI^ cells. The circular plot depicts fold change (sg*MTERF1*/sg*AAVS1*) for each mitochondrial gene across the mtDNA map, with the outer and inner rings corresponding to 95% mtDNA^ΔScaI^and 55% mtDNA^ΔScaI^ cells, respectively. Dashed line indicates the ScaI deletion breakpoints. **(B)** RT-qPCR analysis of relative 7S RNA abundance in sg*AAVS1*- or sg*MTERF1*-targeted mtDNA^ΔScaI^ 95% cells measured at day 12 and day 24 post-nucleofection. Data are shown as mean ± SEM from n = 3 biological replicates. **(C**) Western blot analysis of OXPHOS complex subunits (UQCRC2, COXII, SDHB, and NDUFB8) and GAPDH in 95% and 55% mtDNA^ΔScaI^ cells with sg*AAVS1* or sg*MTERF1* knockout, collected at day 12 and day 24 post-nucleofection in glucose, and at day 24 following a galactose switch at day 12. **(D)** Quantification of relative OXPHOS protein abundance from western blots as in (A) for 95% mtDNA^ΔScaI^ cells. Values are normalized to GAPDH and expressed relative to sgAAVS1 controls. Data are shown as mean ± SEM (n = 3 biological replicates). **(E)** OCR measurements by Seahorse extracellular flux analysis in sgMTERF1-targeted 95% mtDNA^ΔScaI^ cells reconstituted with wild-type *MTERF1*, DNA-binding–deficient *MTERF1* mutants (R162A or 4A), or GFP control, assessed at day 12 post-nucleofection. Cells were cultured in 25 mM glucose (left) prior to Seahorse assay. Data are shown as mean ± SD (n = 4 technical replicates). **(F)** mtFiber-seq analysis of MTERF1 binding site accessibility in wild-type vs mtDNA^ΔScaI^ genomes. (Top) Metaplot of footprint enrichment centered on the MTERF1 binding site. The fraction of reads with a footprint overlapping each position is shown. Smoothing window = 25 bp. Reads were filtered for >10% adenine methylation. (Bottom) Heatmap of footprint occupancy at the MTERF1 binding site for wild-type and mtDNA^ΔScaI^ mtDNA. Each row represents an individual molecule, sorted by decreasing accessibility. Accessible regions are colored dark purple and protected regions orange. Reads were filtered >10% adenine methylation. **(G)** Model of *MTERF1*-loss-mediated high heteroplasmy suppression. Loss of *MTERF1* increases mitochondrial transcript abundance, thereby restoring mitochondrial gene expression and cellular fitness despite persistent high heteroplasmy.

To test whether transcriptional inhibition by MTERF1 requires direct DNA binding, we reintroduced wild-type or DNA-binding-deficient MTERF1 variants into 95% mtDNA^*ΔScaI*^ cells expressing sg*MTERF1*. Wild-type MTERF1 fully reversed the buffering phenotype, suppressing oxygen consumption and proliferation in galactose. In contrast, the DNA-binding-deficient mutants, R162A and 4A (R162A/F243A/Y288A/F322A), failed to suppress respiration and conferred only a partial growth defect, potentially attributable to the residual mtDNA binding that these mutants retain *in vitro*^22^(**Fig. 3E and Fig. S4A-D**). Having established that MTERF1 must engage mtDNA directly to rescue OXPHOS, we next asked whether MTERF1 preferentially associates with wild-type over deleted genomes. To resolve the MTERF1 footprint at single-molecule resolution, we applied mtFiber-seq, which uses a non-specific adenine methyltransferase to record protein occupancy onto individual mtDNA molecules^29^. Consistent with previous observations, MTERF1 occupied its canonical binding site downstream of the rRNA genes on wild-type genomes. We also detected the footprint on deleted genomes at comparable levels (**Fig. 3F**), suggesting that MTERF1 binds both genomes indiscriminately and imposes a genome-nonspecific transcriptional constraint.

Taken together, our data support a model in which MTERF1-mediated transcription termination becomes limiting when the dosage of functional genomes is reduced by large-scale deletions (**Fig. 3G**). At low or no heteroplasmy, MTERF1 binds downstream of the rRNA genes to terminate transcription and modulate 7S RNA levels, maintaining balanced OXPHOS output from the predominant wild-type genomes. As heteroplasmy exceeds the pathogenic threshold, the same termination activity constrains output from an already depleted pool of functional templates, tipping cells into respiratory failure. Loss of MTERF1 lifts this constraint without altering heteroplasmy or copy number, thereby broadly increasing OXPHOS transcripts and reducing 7S RNA. This derepression effectively increases respiratory output from the residual wild-type genomes, restoring oxidative phosphorylation.

### MTERF1 loss broadly buffers against mitochondrial genetic defects

The observation that MTERF1 loss restores OXPHOS capacity by derepressing mitochondrial transcription suggests that it might, more broadly, buffer against OXPHOS insufficiency. To test this prediction systematically, we performed a genome-wide CRISPR/Cas9 screen in wild-type and MTERF1-knockout ARPE-19 cells. Four independent clones of *MTERF1*^-/-^ ARPE-19 cells were pooled (**Fig. S5A**) and, alongside control cells, were transduced with the library of sgRNAs, selected with puromycin, and then split into parallel arms: one maintained in glucose and one shifted to galactose, and grown for 14 days before harvest and deep sequencing of sgRNA representations. (**Fig. 4A, Fig. S5B-E**). We reasoned that glucose withdrawal, which forces cells to rely exclusively on oxidative phosphorylation for ATP production, would reveal fitness defects that glycolysis (in glucose-rich media) would otherwise obscure and provide a sensitized background in which to identify genetic vulnerabilities that MTERF1 loss can compensate for.

**Figure 4:**
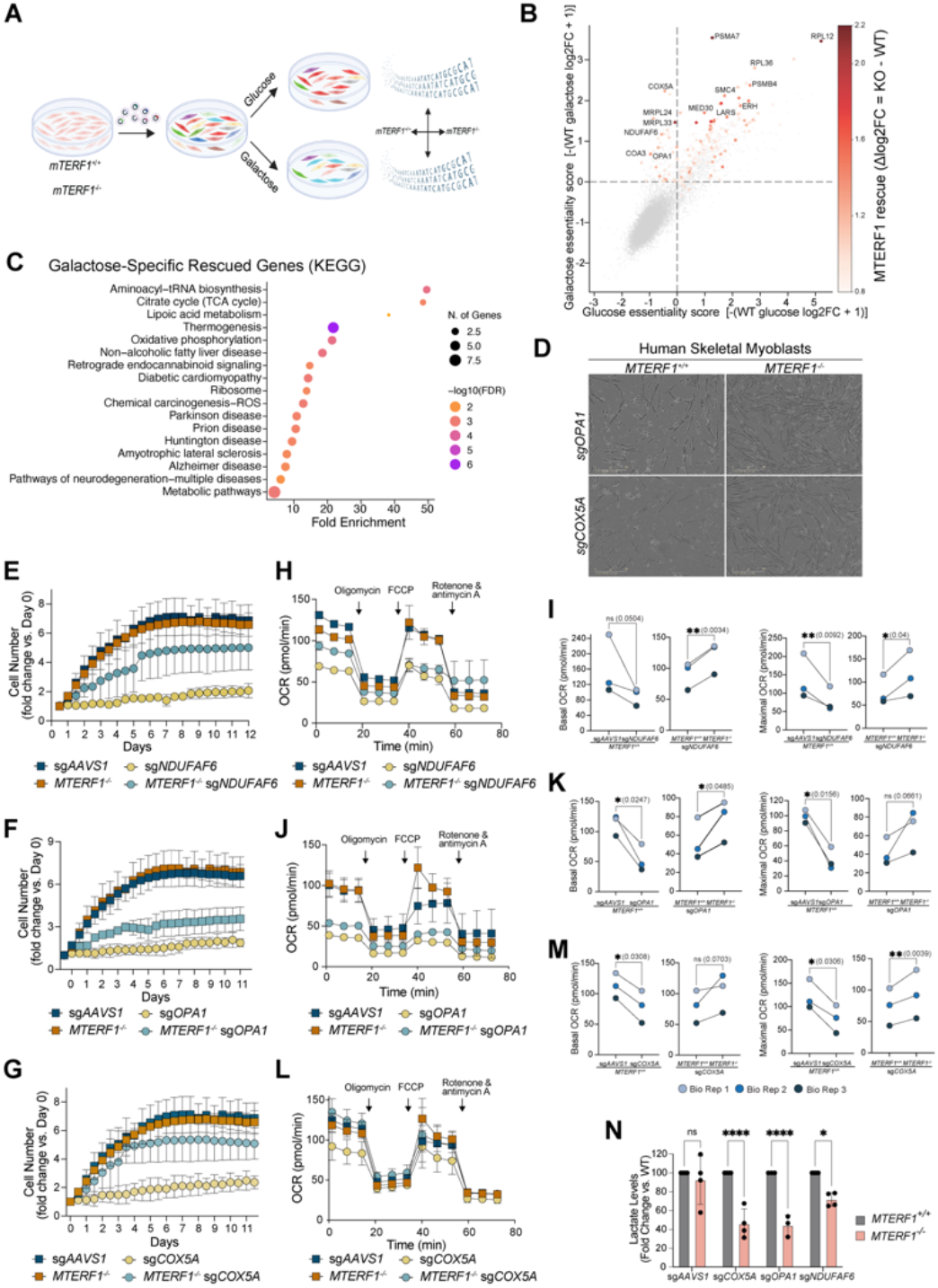
Genome-wide CRISPR screen identifies MTERF1 as a suppressor of multiple metabolic pathways under OXPHOS stress. (A) Schematic of genome-wide CRISPR screen experimental outline. *MTERF1*^+/+^ and MTERF1^-/-^ ARPE19 cells were transduced with TKOv3 sgRNA library, selected with 4 µg/mL puromycin for 48 h beginning 24 h post-infection, and expanded in either glucose or galactose medium for 14 days. (B) The CRISPR/Cas9 screen data were analyzed using the MAGeCK algorithm. Scatter plot represents differential essentiality analysis comparing sg*AAVS1* (WT) and sg*MTERF1* (KO) cells. Each point represents a gene, plotted by glucose essentiality (x-axis; −WT glucose log2FC) versus galactose-specific essentiality (y-axis; −WT galactose log2FC). Points are colored by *MTERF1*-dependent rescue magnitude (Δlog2FC = KO − WT), with selected top hits labeled. Genes in the upper-left quadrant represent OXPHOS-stress dependencies that are alleviated by *MTERF1* loss. (C) KEGG pathway enrichment analysis of galactose-specific genetic vulnerabilities alleviated by *MTERF1* loss, performed using ShinyGO 0.85.1. Dot size indicates gene number; color indicates −log10(FDR). (D) Representative images of sg*COX5A* or sg*OPA1* targeted LHCN-M2 myoblasts (either sg*AAVS1* or sg*MTERF1* genetic backgrounds) at T=8 days in galactose. (E) Growth curves of sg*NDUFAF6* targeted LHCN-M2 myoblasts in sg*AAVS1*- or sg*MTERF1* backgrounds cultured in galactose medium. Data are presented as fold change in cell number relative to day 0. (F) Growth curves of sgOPA1 targeted LHCN-M2 myoblasts in sg*AAVS1*- or sg*MTERF1* backgrounds cultured in galactose medium. Data are presented as fold change in cell number relative to day 0. (G) Growth curves of sgCOX5A targeted LHCN-M2 myoblasts in sg*AAVS1*- or sg*MTERF1* backgrounds cultured in galactose medium. Data are presented as fold change in cell number relative to day 0. (H–M) Oxygen consumption rate (OCR) measured by Seahorse extracellular flux analysis in MTERF1-knockout myoblast lines and control cells cultured in galactose: sg*NDUFAF6* (H and I), sg*OPA1* (J and K), and sg*COX5A* (L and M). Arrows indicate sequential injections of oligomycin, FCCP, and rotenone plus antimycin A. (H, J, and L) show representative Seahorse traces from one biological replicate. (I, K, and M) show mean basal and maximal respiration rates from three biological replicates (see Fig. S6B-C for biological replicates 2 and 3). (N) Mass spectrometry analysis of media lactate levels in sgOPA1, sgCOX5a, and sgNDUFAF6 LHCN-M2 myoblasts under WT or sgMTERF1 genetic backgrounds.

Plotting gene essentiality scores using MAGeCK ^20^ in galactose *vs*. glucose revealed genes that were essential under both carbon sources, as well as a distinct set of genes whose loss was selectively lethal under galactose in *MTERF1*^*+/+*^ cells (**Fig. 4B and Table S5**). Overlaying essentiality scores from *MTERF1*^*-/-*^ cells revealed galactose-specific hits that were substantially rescued upon *MTERF1* loss, including COX5A, MRPL24, MRPS33, NDUFAF6, OPA1, and BCS1L (**Fig. 4B and Table S6**). Pathway enrichment analysis confirmed strong enrichment for aminoacyl-tRNA biosynthesis, the citric acid cycle, and oxidative phosphorylation among the rescued genes (**Fig. 4C and Table S7**). Genes rescued by *MTERF1* loss, regardless of carbon source, were enriched for the proteasome, DNA replication, and ribosome biogenesis pathways (**Fig. S5F**). It is worth noting that loss of *MTERF1* leads to very few new vulnerabilities, including Twinkle helicase (c10orf2) (**Fig. S5G-H)**. Although the top synthetic lethal hits did not map into biological pathways, they might represent biologically meaningful dependencies disrupted by MTERF1 depletion, and characterizing their roles will be a direction for future investigation.

We validated top hits suppressed by *MTERF1* loss, focusing on disease-relevant genes. Specifically, we selected three non-essential genes that each cause inherited mitochondrial disease yet disrupt the respiratory chain in fundamentally different ways. *NDUFAF6*, an assembly factor required to build complex I, and its loss causes Leigh syndrome, a severe infantile neurodegenerative and myopathic disorder^30^. OPA1 is a fusion GTPase that shapes cristae structure and maintains mtDNA; its mutation underlies autosomal dominant optic atrophy and multisystem mtDNA deletion disorders that frequently include mitochondrial myopathy^16,17,31,32^. Lastly, COX5A is a structural subunit of complex IV, and its mutation underlies complex IV deficiency^18,19^. Because skeletal muscle is among the tissues most severely affected in these conditions, we generated single CRISPR/Cas9 knockouts of each gene in human skeletal myoblasts (**Fig. S6A**), which were either wild-type or MTERF1 knockout, and assayed fitness in galactose to enforce OXPHOS dependence.

By phase-contrast imaging, deletion of OPA1 and Cox5A reduced culture density in the MTERF1^+/+^ myoblasts, and concurrent deletion of MTERF1 visibly improved density (**Fig. 4D**). This pattern was mirrored in proliferation assays. Knockout of all three genes impaired myoblast growth over the time course, and *MTERF1* co-deletion increased proliferation relative to the corresponding MTERF1-wild-type knockouts (**Fig. 4E–G**). Respiration followed similarly, with each knockout lowering basal and maximal oxygen consumption by respirometry in the sg*AAVS1* background (**Fig. 4H–M, Fig. S6B-C**), but these reductions were no longer significant when *MTERF1* was concurrently deleted (**Fig. 4I, K, M**). Consistent with a shift away from glycolytic metabolism, extracellular lactate was significantly lower in MTERF1-knockout cells across the *COX5A, OPA1*, and *NDUFAF6* backgrounds (**Fig. 4N**), with reductions reaching significance for each gene knockout but not for the baseline control comparison. Together, these observations suggest that MTERF1 loss can at least partially buffer the fitness and respiratory consequences of functionally distinct mitochondrial insults in skeletal myoblasts, consistent with a broader role for MTERF1 as a modifier of mitochondrial genetic deficiency.

## Discussion

Using a genome-wide CRISPR/Cas9 screen in an isogenic panel of mtDNA^ΔScaI^ cells, we identified *MTERF1* as a nuclear suppressor of mtDNA deletion burden under respiratory-dependent growth. Loss of *MTERF1* restored respiration and proliferation in cells above the >75% pathogenic threshold without altering heteroplasmy or mtDNA copy number. Instead, MTERF1 depletion reduced 7S RNA, raised steady-state mitochondrial transcript abundance, and increased both mtDNA- and nuclear-encoded OXPHOS subunit levels, thereby extracting more functional capacity from the residual pool of wild-type genomes. This rescue extended beyond mtDNA deletions, where a counter-screen showed that *MTERF1* depletion also buffered cells against the loss of diverse nuclear-encoded metabolic genes. Together, these findings position MTERF1 as a context-specific brake on mitochondrial transcriptional reserve, one whose release matters precisely when that reserve becomes limiting.

### A transcript-level means of dosage compensation

Our study suggests that mtDNA deletion disorders are, at their core, a problem of insufficient dosage of functional OXPHOS gene products. Wild-type genomes persist and continue to be transcribed, but their absolute output falls below the level needed to sustain respiration (**Fig 3G**). The majority of therapeutic efforts to date have aimed to fix this by altering the abundance of the mutant genome itself, primarily by eliminating deletion-bearing molecules to shift heteroplasmy ^12,33^. Our results offer a fundamentally different solution. Without altering genome composition or abundance, simply increasing transcriptional output from the existing wild-type templates was sufficient to buffer the deletion and lift cells back above the bioenergetic threshold. This suggests that the deletion phenotype is a constraint on transcript dosage and implies that any intervention that increases the functional transcript pool, such as increasing copy number^34,35^, derepressing transcription, or otherwise boosting mitochondrial gene expression, may produce a comparable benefit. In effect, our data suggest that modest gains in mitochondrial output translate into substantial phenotypic rescue when the underlying reserve is critically low.

An outstanding question concerns the primary physiological function of MTERF1, which remains unsettled. The protein was originally proposed to terminate heavy-strand transcription downstream of the ribosomal genes and to couple termination to reinitiation, thereby defining a transcriptional unit that recycles POLRMT to favor excess rRNA synthesis over the rest of the genome^24^. Subsequent *in vitro* work, however, indicated that termination is directional^25,26^; MTERF1 fully terminates light-strand transcription, preventing run-off interference at rRNA and D-loop. The phenotypes we observe, including decreased 7S RNA, a global rise in mitochondrial transcripts, and notably, a decrease in rRNAs, are consistent with each of these proposed mechanisms.

Beyond relieving essentialities, our genome-wide counter-screen also revealed new genetic vulnerabilities specific to the sg*MTERF1* background (**Fig. S4F**), which may point to additional MTERF1 functions that remain to be uncovered. Most striking is the dependence on the mitochondrial helicase Twinkle, consistent with the reported role of MTERF1 as a contrahelicase that averts replication–transcription collisions at the rRNA locus^25,26^.

### MTERF1 depletion leads to broad buffering against other metabolic deficiencies

The counter-screen in *MTERF1*^*-/-*^ cells identified many nuclear genes whose essentiality was alleviated upon MTERF1 loss, suggesting that the transcriptional buffering mechanism extends beyond mtDNA deletions to a broader range of mitochondrial and metabolic defects. We focused on three linked to human mitochondrial disease that represent distinct points of respiratory-chain failure: *COX5A*, a complex IV subunit mutated in complex IV deficiency^18,19^; *NDUFAF6*, a complex I assembly factor mutated in Leigh syndrome^30^; and *OPA1*, an inner-membrane fusion GTPase mutated in autosomal dominant optic atrophy and linked to mtDNA depletion^16,17,31,32^. A single perturbation rescues lesions within these functionally distinct factors, points to a shared, mutation-agnostic mechanism rather than gene-specific correction. The simplest interpretation is that across this range of partial respiratory-chain defects, pushing more OXPHOS capacity out of the mitochondrial genome is sufficient to compensate for mitochondrial deficiency. Many pathogenic mtDNA mutations, deletions, and nuclear mitochondrial gene defects retain partial activity rather than abolishing it entirely; a mutation-agnostic boost in transcriptional output could push such cells back over the threshold required for respiration. Notably, the rescue of OPA1 loss may operate through an additional, complementary mechanism that further enhances its effectiveness. OPA1 mutations disrupt inner-membrane fusion, cristae remodeling, and mtDNA maintenance, collectively leading to progressive mtDNA depletion and accumulation of multiple deletions in patient tissues^32^. The resulting reduction in mtDNA copy number means that fewer templates are available for transcription, compounding the bioenergetic deficit beyond what would be expected from respiratory-chain dysfunction alone. In this context, elevating transcriptional output per remaining mtDNA copies by releasing the MTERF1-imposed transcriptional block may be particularly effective, compensating for diminished template availability.

### Releasing mitochondrial transcriptional reserve as a therapeutic strategy

Our findings indicate that the wild-type mitochondrial genome holds a transcriptional reserve that, when released, can compensate for a deficit in functional gene dosage. Because this acts on the residual wild-type template rather than on a specific lesion, it offers a mutation-agnostic strategy applicable to disorders ranging from mtDNA deletions and depletion syndromes to nuclear-encoded respiratory chain defects. However, this logic has a clear limit: increasing transcription benefits only loss-of-function mutations. Instead, for gain-of-function, the same increase would amplify the pathogenic species, and *MTERF1* loss would be expected to worsen the phenotype. MTERF1 is tractable because its loss is well tolerated; knockout mice are viable, and normal *MTERF1*^-/-^ cells show no metabolic defect, and the gene is non-essential in DepMap. Its dispensability suggests a therapeutic window in which its inhibition could increase mitochondrial output without compromising baseline

function. MTERF1 is likely only one of several constraints on mitochondrial transcription, and identifying others may broaden this strategy. For mitochondrial disorders that remain without disease-modifying therapy, enhancing output from a patient’s existing wild-type genomes offers a new therapeutic direction.

## ACKNOWLEDGEMENT

We thank Alexandra Schmitt (Electron Microscopy Facility, University of Lausanne, Switzerland) for electron microscopy sample processing and imaging. We acknowledge the Sfeir lab members who commented on the manuscript. This work is supported by grants from the NIH/NIA (R01AG085782-01) and the CHAMP Foundation for A.S, and the Swiss National Science Foundation (310030_21573) for S.M.R.S.I. is supported by the NIH/NIGMS (R35GM162377) and a Karen Toffler Charitable Trust Scholar Award. We acknowledge the use of the Integrated Genomics Operation Core, funded by the NCI Cancer Center Support Grant (CCSG, P30 CA08748). T.K. is the Timmerman Traverse Fellow of the Damon Runyon Cancer Research Foundation (DRG2521-24).

## AUTHOR CONTRIBUTION

T.K. and A.S. conceived the experimental design and implemented the study. T.K and E.C. performed experiments with help from R.C, S.Z, J.L, and S.I. S.Z analyzes sequencing data. J.L in the lab of S.M performed all imaging experiments. R.C in the lab of C.T performed Seahorse assays in Fig 1F, Fig 2A-F. S.I performed Fiber-seq assay. A.S. and T.K. wrote the manuscript with feedback from all authors.

## METHODS

### Cell lines and cell culture

Immortalized Human Skeletal Myoblasts (LHCN-M2) were purchased from ABM (Cat. No T0756) and maintained in PriGrow III (TM003) and PriGrow I (TM001) medium mix (1:4) supplemented with 15% HI-FBS (F4135), 1.4 µg/mL vitamin B12 (Sigma Cas No. 68-19-9), 0.055 µg/mL dexamethasone (Sigma D4902), 0.03 µg/mL zinc sulfate heptahydrate (Sigma Z4750), 0.02 M HEPES (TM058), 10 ng/mL FGF2 (Z101455), 2.5 ng/mL human hepatocyte growth factor (rhHGF) (Z102785) and 1% Penicillin/Streptomycin Solution (G255) and grown in a 37.0°C humidified incubator with 5% CO_2_. Previously published^15^ clones #21 and 23, containing heterogeneous distributions of heteroplasmy, were further single-cell cloned to isolate H87 and H91 from #21 and H95 and H100 from #23. Rho0 cells were isolated from a previously published pool of mito-EJ expressing ARPE-19 cells^15^ and lack of mtDNA confirmed by PCR. Cells with mtDNA deletion were cultured in tetracycline-free glucose medium, consisting of DMEM (Corning MT10027CV) supplemented with 10% tetracycline-free fetal bovine serum (Corning MT35075CV), L-glutamine (2 mM), 1% Penicillin/Streptomycin Solution (G255), 1x MEM non-essential amino acids (Gibco 11140050), and 50 µg/mL uridine. For experiments involving metabolic rewiring, cells were initially seeded in tetracycline-free glucose medium and, on the following day, rinsed with PBS and transferred to Galactose media formulated similarly, with the exception of using glucose-free DMEM (Gibco 11966025), 10% dialyzed FBS (Gibco 26400044), 10 mM Galactose (Sigma G0750), and 1 mM Sodium Pyruvate (Gibco 11360070).

### CRISPR/Cas9 genome-wide synthetic lethal screen

CRISPR/Cas9 screens were performed as described^36^. Briefly, cells were transduced with the lentiviral TKOv3 library at a low MOI (∼0.2–0.3) and selected with puromycin for 48h post-transduction. This was considered the initial time point (T=0). Fig. 1B cells with different genetic backgrounds were selected with 5 ug/ml puromycin and grown for an additional 12 doublings. At T=12, cells were switched into galactose media for 3 days (T=15). The surviving fraction was then enriched in glucose medium for an additional 3 days (T=18) before harvesting the cell pellets. Fig. 4A cells were selected with 4 ug/ml puromycin and grown for 14 doublings in glucose or galactose before harvesting the cell pellets (T=14). Cell pellets were frozen at each time point for genomic DNA (gDNA) isolation. A library coverage of ≥500 cells per sgRNA was maintained at every step. gDNA from cell pellets was isolated using the Quick-DNA Midiprep Plus Kit (ZymoResearch, D4075) and genome-integrated sgRNA sequences were amplified by PCR 25 using the Q5 Mastermix (New England Biolabs Next UltraII). i5 and i7 multiplexing barcodes were added in a second round of PCR, and final gel-purified products were sequenced on Illumina HiSeq2500 or NextSeq500 systems to determine sgRNA representation in each sample.

MAGeCK gene-level LFC rankings between T=12 and T=18 were used for plotting Figure 1C. Guides with fewer than 50 normalized reads at Day 0 or zero reads at Day 12 were excluded. Normalized sgRNA read counts generated by MAGeCK^20^ were used to calculate sgRNA-level log2 fold changes between T=0 and T=14 conditions in Fig 4B. Guides with fewer than 50 normalized reads at Day 0 or zero reads at Day 14 were excluded. Gene-level log2 fold changes were calculated as the mean log2 fold change across sgRNAs targeting each gene. Alleviated genes were defined as those with a mean LFC < −1 in the sgAAVS1 background, a mean LFC ≥ −1 in the sgMTERF1 background, and a differential mean LFC (sgMTERF1 − sgAAVS1) > 0.8. Synthetic lethal genes were defined as the inverse: mean LFC < −1 in sgMTERF1, mean LFC ≥ −1 in sgAAVS1, and differential mean LFC < −0.8. Library coverage (mean reads per sgRNA), sequencing depth (total mapped reads), and read count distributions were calculated from normalized sgRNA count tables at T=0 and T=12 timepoints for Fig. 1B, and T=0 and T=14 timepoints for Fig. 4A. Read count distributions were visualized with a 95th percentile cutoff to exclude outliers.

Each screen performance was assessed using BAGEL2 (Bayesian Analysis of Gene Essentiality with Location)^37,38^ by comparing the log_2_ fold change (LFC) distributions of reference essential and non-essential gene sets. In Fig. S1G, LFC values were calculated between T=0 and T=12 in glucose timepoints for sgRNAs targeting known essential genes and non-essential genes^36^.In Fig. S5E, they were calculated between T=0 and T=14. Kernel density estimation (KDE) was used to visualize the separation between distributions. A quality score was calculated as the absolute difference between the median LFC of essential and non-essential gene sets (S1G quality score = 1.81; S5E quality score = 1.79), indicating clear separation between essential and non-essential guides and successful screening performance. Analyses were performed using BAGEL2 and Python (scipy, numpy, matplotlib).

### Plasmids

MTERF1 expressing plasmid was sourced from Addgene (TFORF1832 (Plasmid #143160). R162A and 4A construct gblocks were designed and ordered from Integrated DNA Technologies (IDT). DNA constructs were generated using the NEBuilder® HiFi DNA Assembly Kit (New England Biolabs) according to the manufacturer’s guidelines. Briefly, vector backbones and insert fragments with overlapping homology arms were assembled in vitro, and assembled products were introduced into chemically competent E. coli by heat-shock transformation. Transformed cells were allowed to recover in SOC medium and plated onto selective agar plates. Colonies were grown overnight at 37 °C, and plasmids were isolated and verified by Plasmidsaurus ZeroPrep sequencing prior to downstream applications.

### Lentiviral Production and Transduction

Lentivirus were produced in HEK293T cells by transient transfection using a polyethylenimine (PEI)–based method. Packaging plasmids and the lentiviral transfer vector encoding the gene of interest were combined in saline solution and complexed with PEI prior to addition to cells. The transfection mixture was added dropwise to HEK293T cells, which were incubated overnight before replacement with fresh culture medium. Viral supernatant was collected 24 h after medium change, clarified by filtration through a 0.45 µm filter, and either used immediately or stored at −80 °C until use. Target cells were transduced with viral supernatant under standard culture conditions and selected as indicated for downstream experiments.

### Genomic DNA Purification

DNA was purified for mtDNA deletion detection, measurements of heteroplasmy and copy number by qPCR. Cell pellets harvested were resuspended in 500 µl PBS buffer containing 0.2% (w/v) SDS, 5 mM EDTA, and 0.2 mg/ml Proteinase K and incubated at 50°C for 3-4 hours with constant shaking at 1,000 rpm. DNA was precipitated by adding 0.3 M Sodium Acetate, pH 5.2, and 600 µl isopropanol and kept at -20°C for 2+ hours. Precipitated DNA was centrifuged at 21,000 x g at 4°C for 15 min, followed by a cold 70% ethanol wash. DNA was then resuspended in TE buffer (10 mM Tris-HCl, pH8.0, 0.1 mM EDTA) and quantified by Nanodrop.

### Nucleofection

CRISPR–Cas9 ribonucleoprotein (RNP) complexes were assembled by incubating recombinant Cas9 nuclease with sgRNA at room temperature for 15–30 min. Cells were harvested and resuspended in nucleofection buffer (Lonza P3 buffer with supplement), and 5 × 10^5^ cells per reaction were mixed directly with pre-assembled RNP complexes. Nucleofection was performed using a Lonza 4D-Nucleofector system, following the manufacturer’s recommendations using code CA137. Cells were recovered with RPMI for 15 minutes and plated overnight before changing media 24 hours later. sgMTERF1 experiments Fig. 1-3 were performed starting at T=12 days post nucleofection. sgMTERF1 experiments in Fig. 4 in both ARPE-19 and LHCN-M2 Myoblasts were performed in individually validated and pooled clones generated post nucleofection.

### Quantitative PCR

Mitochondrial DNA heteroplasmy was quantified by quantitative PCR (qPCR) using total genomic DNA as input as previously described^15^. Briefly, mtDNA deletion junction and adjacent wild-type sequence alongside nuclear reference locus (B2M) were independently amplified using PowerUp^−^ SYBR^−^ Green Master Mix (Applied Biosystems) and target-specific primers. qPCR reactions were performed in 10 µl volumes with 250 nM primers and 2 ng genomic DNA per reaction using standard cycling conditions on Applied Biosystems QuantStudio 6 Real-Time PCR System. Relative abundance was calculated from threshold cycle (Ct) values. Deletion heteroplasmy was determined by calculating the fraction of deletion-containing mtDNA relative to the combined abundance of deletion-bearing and wild-type mtDNA genomes. mtDNADScaI heteroplasmy = (2^-(CtDScaI) / (2^-(CtDScaI) + (2^-(Ct WT)) ^*^ 100%.

### Total RNA Extraction and RT-qPCR

Total RNA was extracted using the QIAGEN RNeasy Mini Kit according to the manufacturer’s instructions. Complementary DNA (cDNA) was synthesized from purified RNA using the iScript^−^ cDNA Synthesis Kit (Bio-Rad) following the manufacturer’s protocol. Mitochondrial transcripts including RNR1, RNR2, ND1, ND2, ND5, CYTB, ND6, and ATP6 were quantified using target-specific primers designed with the IDT Real-Time qPCR Primer Design Tool. All primer sets were validated for amplification efficiency and specificity prior to use.

Quantitative PCR reactions were performed in 10 µl volumes with 250 nM primers using PowerUp^−^ SYBR^−^ Green Master Mix (Applied Biosystems). cDNA was diluted 1:100 before amplification, and reactions were run on a QuantStudio 6 Real-Time PCR System using standard cycling conditions recommended by the manufacturer. Transcript abundance was quantified based on threshold cycle (Ct) values and normalized to B2M expression.

### RNA-Sequencing and Analysis

Cell pellets were resuspended in DNA/RNA Shield (Zymo R1100-250), and RNA sequencing was performed using the Plasmidsaurus DGE RNA-seq platform. Sequencing reads were aligned by Plasmidsaurus using the GRCh38 reference genome (Ensembl release 114). Gene-level read counts were quantified using featureCounts v2.1.1 from the Subread package using the Ensembl release 114 GTF annotation. An additional annotation feature corresponding to the mitochondrial 7S RNA locus (chrM:260–382) was added to the annotation file prior to counting. Gene counts were generated from deduplicated aligned BAM files produced by the Plasmidsaurus DGE RNA-seq pipeline. Differential expression analysis was performed using DESeq2. Genes with an adjusted P value < 0.05 and an absolute log2 fold change > 0.5 were considered significantly differentially expressed. Volcano plots were generated using DESeq2 log2 fold changes and adjusted P values. Heatmaps were generated from DESeq2-normalized gene counts.

### Western blot

Cells were harvested and lysed under conditions optimized for detection of mitochondrial and OXPHOS proteins. Cells were lysed in RIPA buffer supplemented with detergents and buffered salts, followed by sonication at 4 °C. Protein concentration was determined using a bicinchoninic acid (BCA) assay, and equal amounts of total protein were mixed with Invitrogen NuPAGE^−^ LDS Sample Buffer (4X) (Cat NP0007). Lysates were mixed and denatured by heating at 95 °C for 10 min. Equal volume were loaded per onto Invitrogen Bis-Tris Protein Gels, 4–12% (Cat NP0321BOX). Proteins were transferred to nitrocellulose membranes using semi-dry Turbo transfer systems (Bio-Rad).

Membranes were blocked in 5% non-fat milk in TBST and incubated with primary antibodies diluted in blocking buffer overnight at 4 °C. Following washes in TBST, membranes were incubated with Licor IRDye® Secondary Antibodies and washed in TBST. Fluorescence signals were acquired using a Licor Odyssey CLx Imager. Protein abundance was quantified using ImageStudio software, with GAPDH serving as a loading control.

Primary antibodies used in this study targeted mitochondrially encoded and nuclear-encoded OXPHOS subunits, and loading controls, including COXII, NDUFB8, UQCRC2, ATP5A, SDHB, GAPDH, (sources listed in below). Abcam (Cat ab110411) Total OXPHOS Human WB Antibody Cocktail (1:1000) CellSignaling (2118) GAPDH (14C10) Rabbit Monoclonal Antibody

### Proliferation assay (IncuCyte)

Cells were harvested, washed, and resuspended in fresh glucose or galactose medium. Cells were seeded into 48-well plates at 5 × 10^3^ cells per well in 250 µl medium per technical replicate. Plates were placed in the IncuCyte and imaging was initiated on the day of seeding. Culture medium was replenished every three days, and cell growth was monitored continuously throughout the experiment. Confluence measurements were normalized to day 0, and relative fold changes in confluence were used to compare growth rates across conditions. Plates were imaged every 12 hours on the Incucyte S3 Live-cell analysis system (Sartorius) using brightfield and 10x objective. 16 fields per well were captured. Cell confluency was measured using Incucyte analysis software with settings as follows: 0.5 Segmentation Adjustment, 700 µm2 Hole Fill, 2000 µm2 minimal area.

### Oxygen consumption measurements

Mitochondrial respiration was assessed by measuring oxygen consumption rate (OCR) using a Seahorse XFe96 Extracellular Flux Analyzer (Agilent), according to the manufacturer’s guidelines. Cells were seeded into Seahorse XF microplates at a density of 3 × 10^4^ cells per well and allowed to adhere overnight. Prior to the assay, growth medium was replaced with Seahorse assay medium (Agilent DMEM) supplemented with glucose or galactose, and plates were equilibrated under non-CO_2_ conditions. OCR was recorded under basal conditions and following sequential injection of oligomycin, FCCP, and a rotenone/antimycin A mixture to assess mitochondrial respiratory parameters as recommended by the manufacturer. Cells in well were pooled and counted after each experiment to normalize raw OCR data.

### Lactate analysis using GC-MS

Cells were seeded in regular glucose media at 10% density and grown for 5 days before harvesting 1 ml of media. 20 µL of medium was collected and quenched with 1 mL of ice-cold 80% methanol. Samples were then centrifuged at 21,000g for 30 minutes to pellet protein debris from serum-containing medium. The resulting supernatant was dried for 6 hours using a vacuum concentrator (Genevac EZ-2 Elite). Dried metabolites were resuspended in 20 mg/mL methoxyamine hydrochloride (Sigma, 226904) prepared in pyridine (Thermo Fisher Scientific, TS-27530) and incubated at 30°C for 90 minutes. Samples were subsequently derivatized with MSTFA containing 1% TMCS (Thermo Fisher Scientific, TS-48915) at 37°C for 30 minutes. GC-MS analysis was carried out using an Agilent 7890A gas chromatograph coupled to an Agilent 5975C Mass Selective Detector with electron impact ionization.

Samples were analyzed in splitless mode with helium as the carrier gas at a constant flow rate of 1 mL/min. One microliter of each derivatized sample was injected onto an HP-5MS column (15 m × 0.25 mm, 0.25 µm film thickness). The inlet temperature was maintained at 250°C, and the oven temperature was increased from 60°C to 290°C over 25 minutes. Data analysis was performed using MassHunter Quantitative Analysis software (v.10.0, Agilent Technologies). Lactate was quantified using the ion at m/z 219. Chromatographic peaks were manually inspected and confirmed by comparison with reference spectra.

### Live Imaging

For live-cell fluorescence imaging, cells were seeded in glass-bottomed cell culture dishes (ibidi, 81158) and cultured in their normal medium (without phenol red) at 37°C, 5% CO_2_. Prior to imaging, cells were stained with 1:10’000 dilution of SYBR Gold (Thermo Fisher Scientific, S11494) and 50 nM TMRE for 10 min, followed by two washes with media and imaging immediately in the transparent media. Live imaging was performed in a Nikon AX-R laser scanning confocal microscope equipped with the NSPARC detector, using a Nikon Plan Apochromat 60X 1.42 NA oil immersion objective and processed on-the-fly in the Nikon NIS-Elements software. Mitochondrial network and nucleoid morphology quantification were performed using built-in CellProfiler 4.2.5 functions. Briefly, the mitochondrial signal was segmented using Otsu thresholding (via the IdentifyPrimaryObjects module), which was then used as a mask for nucleoid detection with the same module. Then, the segmented objects were measured by the MeasureObjectSizeShape module. The aspect ratio of mitochondria was calculated as the ratio between the major and minor axis lengths (AreaShape_MajorAxisLength/AreaShape_MinorAxisLe ngth). For electron microscopy, cells were seeded in glass-bottomed cell culture dishes (Mattek P35G-1.5-14-C).

### Electron Microscopy

Electron microscopy experiments were performed at the Electron Microscopy Facility of the Faculty of Biology and Medicine of Université de Lausanne. Cells were seeded in glass-bottomed cell culture dishes (Mattek P35G-1.5-14-C) and fixed for 1h with 2.5% glutaraldehyde (EMS) in 0.1M phosphate buffer (pH 7.4). They were then post-fixed for 1h with 2.5% glutaraldehyde, 1% osmium tetroxide (EMS) and 1.5% potassium ferrocyanide (Sigma) in 0.1M phosphate buffer (pH 7.4). Cells were rinsed in distilled water, then dehydrated in a graded ethanol (Sigma) series : 30%, 70%, and 3 x 100% for 10min each. This was followed by infiltration in Epon 812 (EMS) overnight. Epon 812 was then replaced by fresh resin, and samples were polymerized for 48h in a 60°C oven. Longitudinal ultrathin sections (60 nm) were collected 1 µm below the surface using a Leica EM UC7 ultramicrotome, on copper 2 x 1mm slot grids (EMS) coated with polyetherimide (Aldrich). Sections were post-stained with 2% uranyl acetate (Sigma) in water and Reynolds lead citrate (Sigma) for 10min each. Micrographs were acquired using a FEI Tecnai Spirit (FEI) at 80kV equipped with a TVIPS F416 camera (TVIPS).

### mtFiber-seq

Hia5 methyltransferase purification, mitochondrial isolation, and mtFiber-seq were performed as previously described^29^. Cells were maintained in DMEM supplemented with Tet-free FBS (TaKaRa 631368), Pen/Strep (Thermo Fisher Scientific 15130122), 0.1 mM NEAA (Thermo Fisher Scientific 11-140-076), 50 µg/mL uridine, and 2 mM L-glutamine. For glucose/galactose treatment, cells were plated at 2x density for galactose conditions. The following day, maintenance media was replaced with either glucose or galactose treatment media. Both media were prepared using DMEM lacking glucose (Thermo Fisher Scientific 11966025) supplemented with dialyzed FBS (Thermo Fisher Scientific A3382001), Pen/Strep, 0.1 mM NEAA, 50 µg/mL uridine, 2 mM L-glutamine, and 1 mM sodium pyruvate (Thermo Fisher Scientific 11360070), with the addition of either 25 mM glucose or 10 mM galactose, respectively. Cells were harvested after 3 days of treatment for mtFiber-seq.

Cells were pelleted by spinning at 150 x g for 5 minutes at 4°C. Media was removed, and cells were resuspended in 4 mL Cell Lysis Buffer (10 mM Tris, pH 7.5; 10 mM NaCl; 1.5 mM MgCl_2_). Cells were allowed to swell on ice for 7.5 minutes, then lysed by douncing in a glass dounce homogenizer with 20 strokes. 2 M Sucrose T10E20 Buffer (10 mM Tris, pH 7.6; 1 mM EDTA, pH 8.0; 2 M Sucrose) was added to bring the final sucrose concentration to 250 mM. Cell debris was pelleted by centrifuging lysates at 1,300 x g for 3 minutes at 4°C. The supernatant was transferred to fresh Eppendorf tubes and spun again at 1,300 x g for 3 minutes at 4°C. The supernatant was transferred to fresh Eppendorf tubes, and mitochondria were pelleted by spinning at 18,000 x g for 15 minutes at 4°C. Mitochondria pellets were resuspended in Permeabilization Buffer (20 mM Tris, pH 7.4; 70 mM Potassium Acetate; 250 mM Sucrose; 0.25% Tween-20; 0.25% Igepal CA-630) and incubated on ice for 10 minutes. Mitochondria were then pelleted by centrifugation at 18,000 x g for 15 minutes at 4°C. Supernatant was removed, and pellets were resuspended in 56 µL mtFiber-seq reaction buffer (20 mM Tris, pH 7.4; 70 mM Potassium Acetate; 250 mM Sucrose). S-adenosylmethionine was added to a final concentration of 0.8 mM and tubes were transferred to a thermocycler prewarmed to 37°C. Reactions were allowed to proceed for 10 minutes at 37°C before quenching with 3 µL 20% SDS. Sample volume was increased to 200 µL by adding reaction buffer and 7 µL 20% SDS. Protein was degraded by adding 2 µL 18.2 mg/mL Proteinase K (Millipore Sigma 3115887001) and incubating at 55°C for 1 hour. Phenol:chloroform:isoamyl alcohol extraction was performed to extract DNA, followed by ethanol precipitation using 0.1 volumes 3 M Sodium Acetate pH 5.5 and 2.5 volumes cold 100% ethanol. DNA was pelleted, washed with 70% ethanol, and air dried for 10 minutes at room temperature before resuspending in 81 µL water. 10 µL NEB CutSmart Buffer was added, samples were treated with 3 µL RNase A (Thermo Fisher Scientific AM2270), 3 µL XmaI (NEB R0180), and 3 µL BamHI-HF (NEB R3136), and incubated at 37°C for 1 hour. A phenol:chloroform:isoamyl alcohol extraction was performed after bringing sample volume up to 200 µL, followed by a chloroform:isoamyl alcohol extraction to remove residual phenol. DNA was precipitated as before, pelleted, washed with 70% ethanol, and air dried. DNA was resuspended in 75 µL Qiagen Buffer EB, quantified using the Qubit hsDNA Kit (Thermo Fisher Scientific Q32851) and sequenced in a PacBio Revio system at the University of Maryland Institute for Genome Sciences, following library preparation according to manufacturer instructions.

### mtFiber-seq analysis

PacBio HiFi reads were generated by the University of Maryland Institute for Genome Sciences with kinetic information retained. Fibertools (v0.8.2) predict-m6a was used on PacBio CCS BAM files to predict m6A modified bases. Data were aligned to a custom reference genome containing wild-type mitochondrial DNA (chrM) and a ΔScaI deletion mtDNA (chrM_del) using pbmm2 (v26.1.99) with the –preset CCS, --sort, --unmapped, and –log-level INFO flags. Fibertools ft extract was used with a minimum score of 252 to extract m6A positions and generate BED files.Footprints were called on aligned, m6A-tagged BAM files using FiberHMM (v2.10.5) with the –mode pacbio-fiber, --circular, and –scores flags and the bundled hia5_pacbio.json model. The –circular flag tiles each read three times prior to HMM inference to avoid boundary artifacts arising from the circular mitochondrial genome. Footprint and methylase-sensitive patch (MSP) coordinates were extracted from the output BAM files using fiberhmm-extract with the –footprint, --msp, and – bed-only flags. Full-length reads were defined as those spanning the complete reference sequence based on coordinates in the m6A BED file. Reads were filtered for accessibility using a threshold of >= 10.8% adenine methylation (>=1,000 m6A sites for chrM; >=786 m6A sites for chrM_del). Footprint heatmaps were generated by expanding BED12 footprint blocks into per-read genomic intervals and sorting reads by decreasing m6A count. Metaplots of footprint enrichment were computed as the fraction of reads with a footprint overlapping each genomic position, smoothed with a 25 bp rolling mean window.

### Pathway / KEGG enrichment analysis

Pathway enrichment analysis was performed using ShinyGO v0.85.1^39,40^. Genes identified as hits from the CRISPR screen were submitted as input with the following parameters: species, *Homo sapiens*; FDR cutoff, 0.05; pathway size minimum/maximum, default settings. Enriched KEGG pathways^41^ were reported.

## Statistical analysis

Statistical analysis was performed using GraphPad Prism (v10). Two-way ANOVA with Šídák’s multiple comparisons test was used to compare sgAAVS1 and sgMTERF1 groups across culture conditions. Fig. S2F was analyzed using ordinary one-way Anova with Tukey all-pairs comparisons. Data are presented as mean ± SD (n=3 biological replicates). P-values < 0.05 were considered statistically significant.

## Data and code availability

Raw sequencing data from the CRISPR screen have been deposited in the NCBI Gene Expression Omnibus (GEO) under accession number (available upon publication). CRISPR screen analysis was performed using MAGeCK^20^ and BAGEL2^38^

## Supplemental Figures and Legends

**Figure S1:**
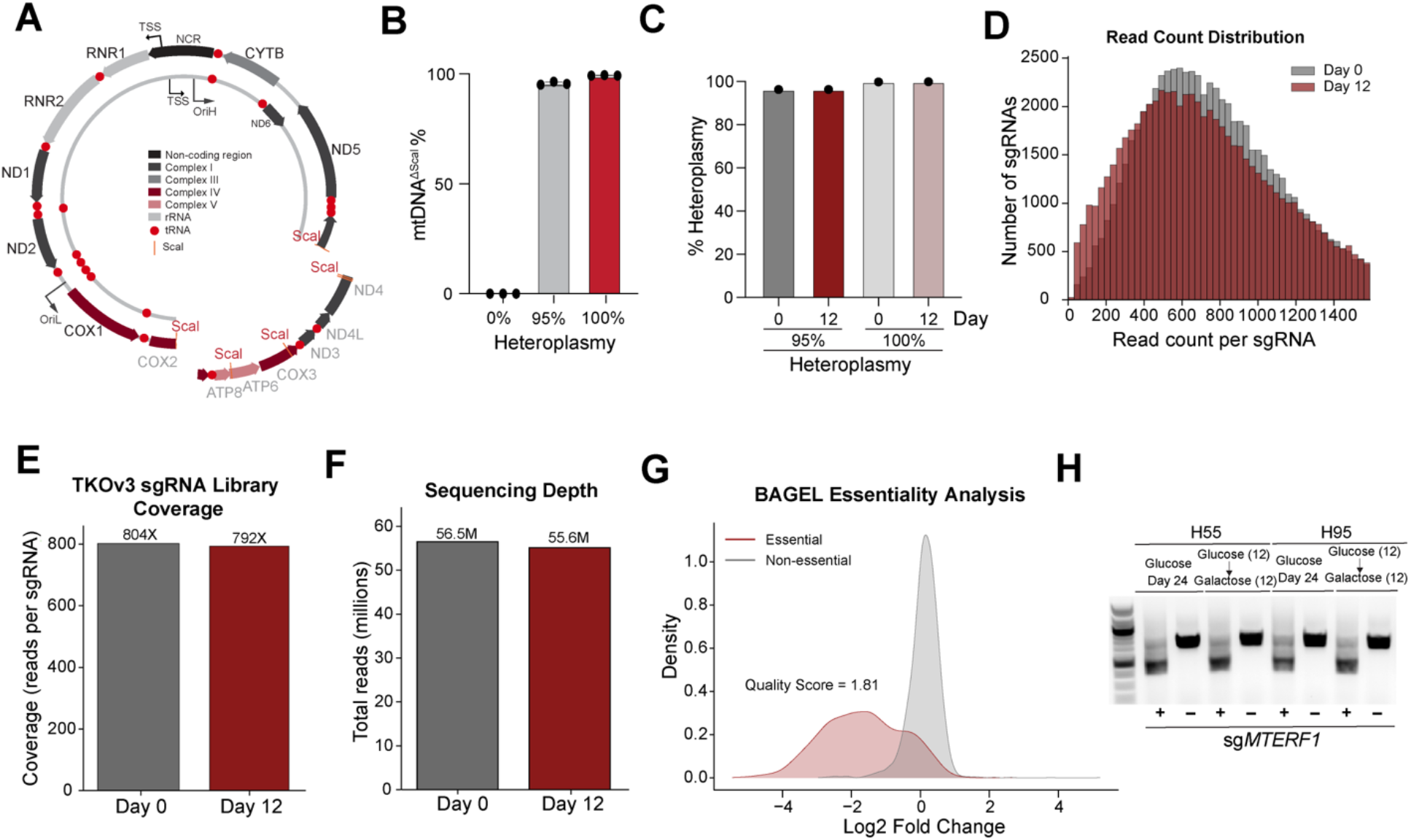
Characterization of heteroplasmic cell lines and genome-wide CRISPR screen quality metrics. **(A)** Schematic of mtDNA containing a large-scale ScaI deletion. Genes lost: COX2, ATP8, ATP6, COX3, ND3, ND4L, ND4, and tRNAs (K, G, & R). **(B)** Quantitative PCR (qPCR) analysis of mtDNA^ΔScaI^ levels of cells in Fig 1A. **(C)** qPCR analysis of mtDNA^ΔScaI^ 95% and 100% clones from the CRISPR screen in Figure 1 at Days 0 and 12. **(D)** Read count distribution at Day 0 vs Day 12 of guides in Figure 1B. **(E)** TKOv3 sgRNA guide library coverage of the guides in Figure 1B **(F)** Sequencing depth of samples in (D). **(G)** BAGEL Essentiality score analysis for screen quality in Figure 1B. **(H)** Agarose gel electrophoresis of PCR-amplified MTERF1 locus in sg*AAVS1*- or sg*MTERF1*-targeted 55% and 95% mtDNA^ΔScaI^ cells.

**Figure S2:**
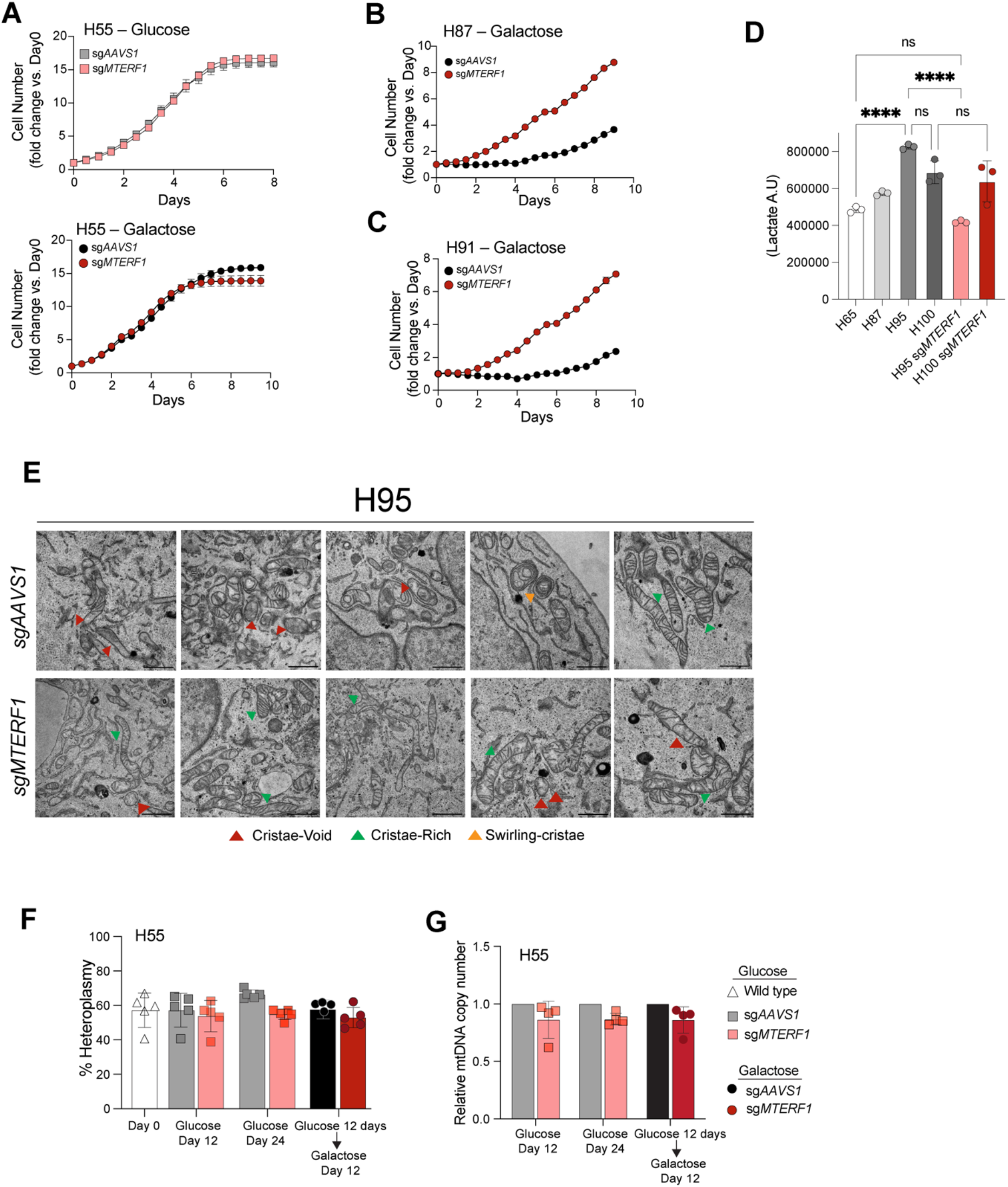
MTERF1 loss enhances proliferation across multiple heteroplasmic cell lines and alters mitochondrial ultrastructure. **(A)** Growth curves of sg*AAVS1*- or sg*MTERF1*-targeted 55% mtDNA^ΔScaI^ cells cultured in glucose (top) or galactose (bottom) medium. Data are presented as fold change in cell number relative to day 0. **(B)** Growth curves of sg*AAVS1*- or sg*MTERF1*-targeted 87% mtDNA^ΔScaI^ cells cultured in galactose medium. Data are presented as fold change in cell number relative to day 0. **(C)** Growth curves of sg*AAVS1*- or sg*MTERF1*-targeted 91% mtDNA^ΔScaI^ cells. Same conditions as in (B). **(D)** Mass spectrometry analysis of media lactate levels in indicated heteroplasmy levels in wild-type or sg*MTERF1* ARPE-19 cells. **(E)** Representative images of electron microscopy from Fig. 2J. **(F-G)** qPCR quantification of heteroplasmy (F) and relative mtDNA copy number (G) in sg*AAVS1*- or sg*MTERF1*-targeted 55% mtDNA^ΔScaI^ cell lines at the same time points as in (Fig 2K-L).

**Figure S3:**
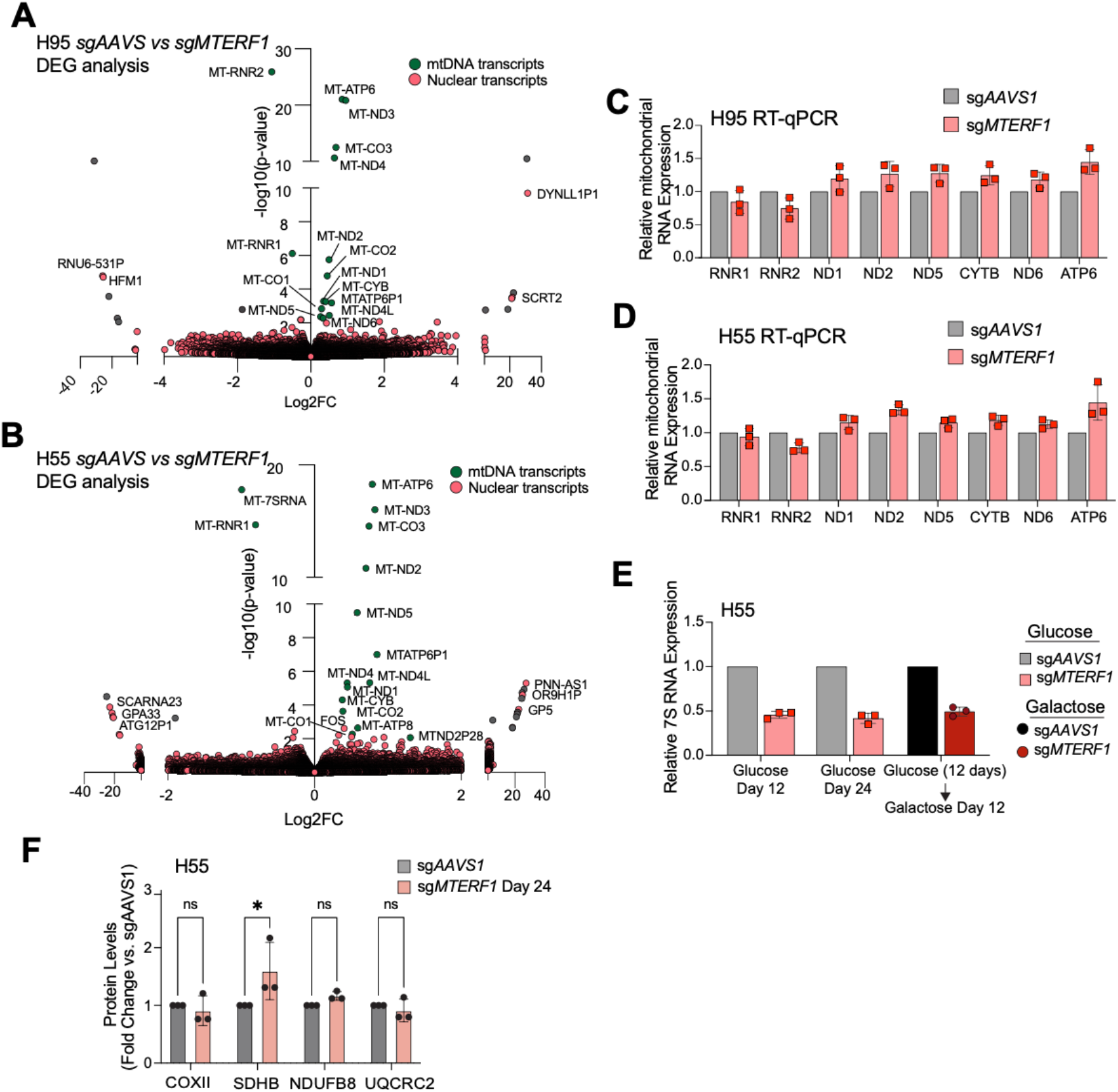
MTERF1 depletion upregulates mitochondrial transcripts without affecting mtDNA copy number or heteroplasmy. **(A)** RNA-sequencing differential gene expression analysis of sgAAVS1 vs sgMTERF1 95% mtDNA^ΔScaI^ cells. mtDNA derived transcripts are shaded in green. Nuclear transcripts are shared in red. Pseudogenes are not labeled and shaded in grey. (**B)** Same analysis as in (A) in 55% mtDNA^ΔScaI^. **(C)** RT-qPCR analysis of mitochondrial DNA transcripts in sg*AAVS1*- or sg*MTERF1*-targeted 95% mtDNA^ΔScaI^ cells. **(D)** RT-qPCR analysis of mitochondrial DNA transcripts in sg*AAVS1*- or sg*MTERF1*-targeted 55% mtDNA^ΔScaI^ cells. **(E)** RT-qPCR analysis of relative 7S RNA abundance in sg*AAVS1*- or sg*MTERF1*-targeted mtDNA^ΔScaI^ 55% cells measured at day 12 and day 24 post-nucleofection. Data are shown as mean ± SEM from n = 3 biological replicates. **(F)** Quantification of Western Blot in Fig. 3C.

**Figure S4:**
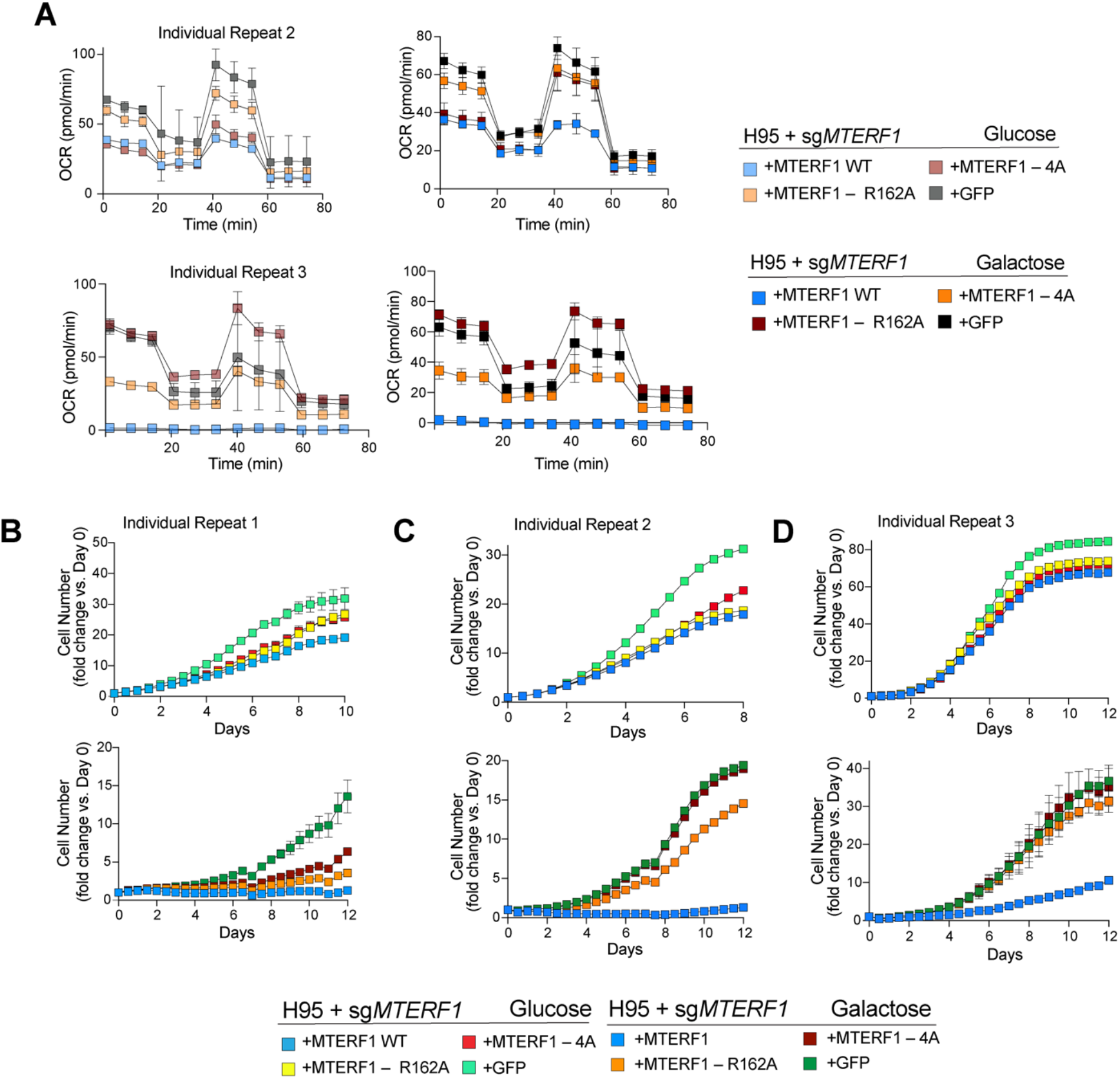
DNA-binding activity of MTERF1 is required to suppress mitochondrial respiration and restrict proliferation in galactose. **(A)** Biological repeats of Fig. 3E. **(B)** Growth curves of sgMTERF1-targeted 95% mtDNA^ΔScaI^ cells reconstituted with wild-type *MTERF1*, DNA-binding–deficient *MTERF1* mutants (R162A or 4A), or GFP control, assessed at day 12 post-nucleofection. Cells were cultured in 25 mM glucose (top) or 10 mM galactose (bottom) medium. Data are shown as mean ± SD (n = 4 technical replicates**). (C)** Biological repeat of (B). **(D)** Biological repeat of (B).

**Figure S5:**
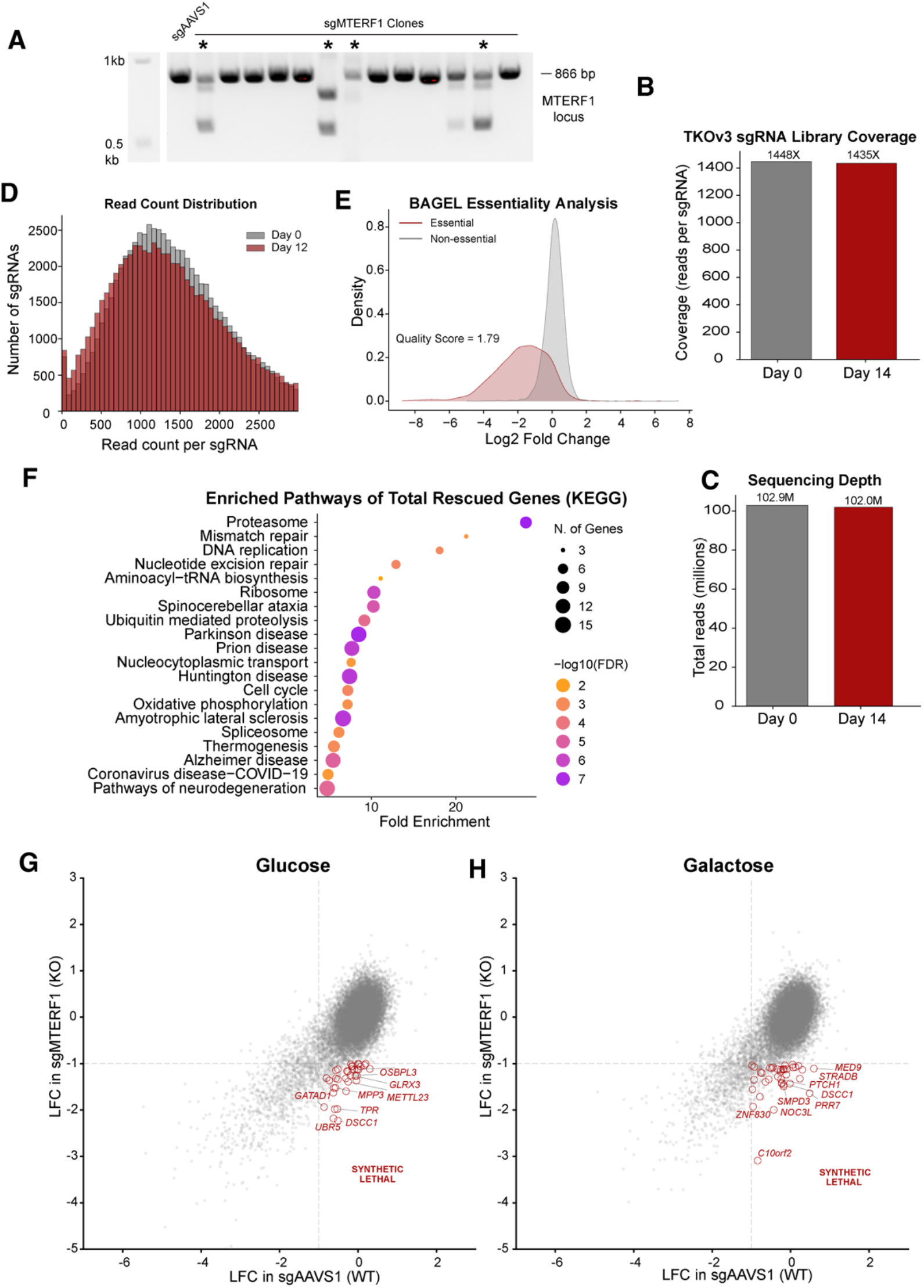
Genome-wide CRISPR/Cas9 screen identifies suppressor and synthetic lethal interactions with MTERF1 loss. **(A)** Agarose gel electrophoresis of PCR amplified MTERF1 locus in sg*AAVS1*- or sg*MTERF1*-targeted ARPE-19 Clones. **(B)** TKOv3 sgRNA guide library coverage of the genome-wide CRISPR screen in wild-type cells from Fig 4A Day 0 vs Day 14 in glucose. **(C)** Sequencing depth of samples in (A). **(D)** Read count distribution at Day 0 vs Day 14 of guides in (A). **(E)** BAGEL Essentiality score analysis for screen quality. (E) Ranked gene essentiality analysis from (D). **(F)** KEGG pathway enrichment analysis of genome-wide genetic vulnerabilities alleviated by *MTERF1* loss, performed using ShinyGO 0.85.1. **(G)** The CRISPR/Cas9 screen data were analyzed using the MAGeCK algorithm. Scatter plot represents differential essentiality analysis comparing sg*AAVS1* (WT) and sg*MTERF1* (KO) cells. Each point represents a gene, plotted by glucose essentiality (x-axis; LFC in sgMTERF1) versus galactose specific essentiality (y-axis; LFC in WT). Genes in the lower-right quadrant represent dependencies under OXPHOS stress that are worsened by *MTERF1* loss. **(H)**. Same analysis as in (G) in galactose.

**Figure S6:**
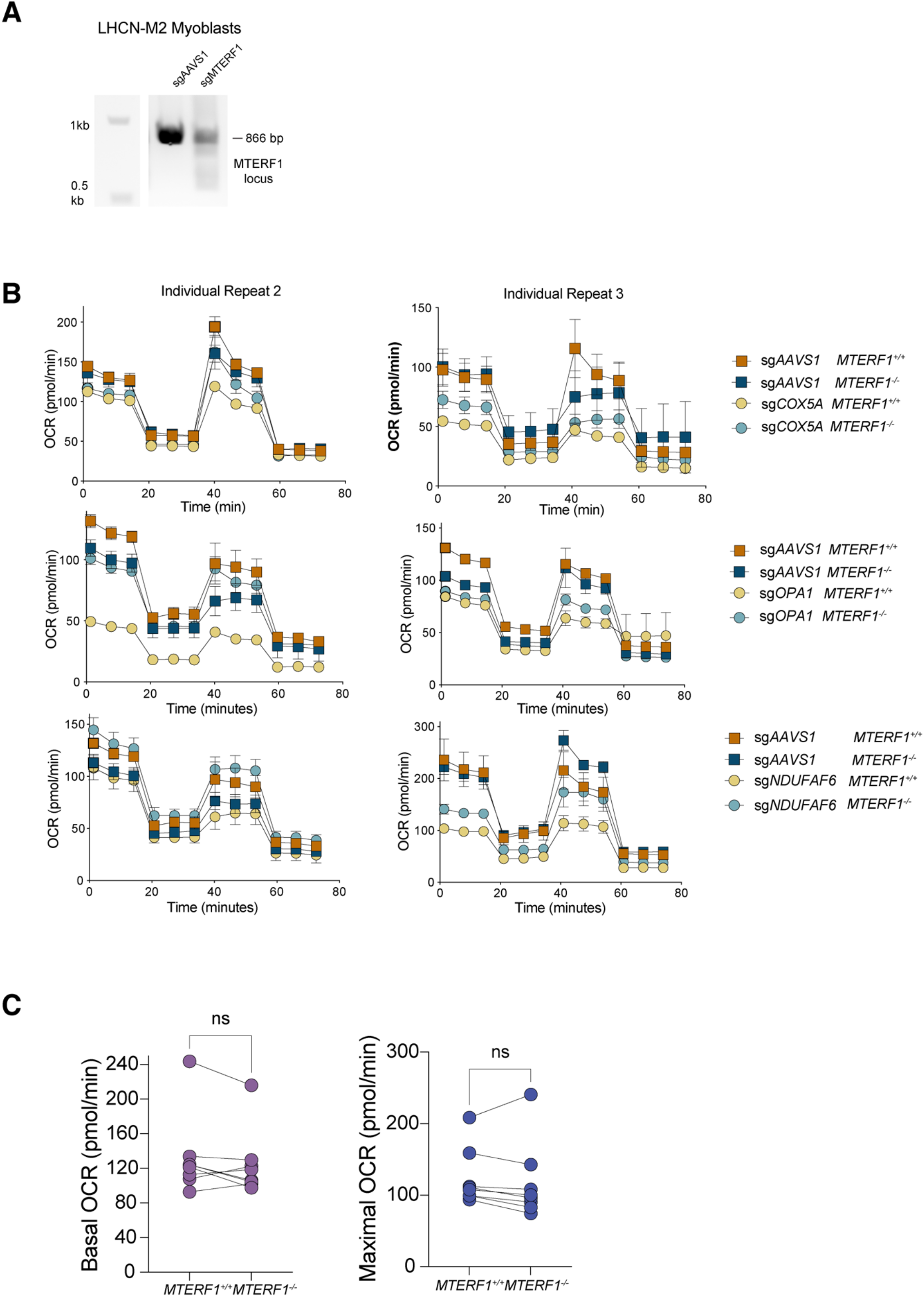
MTERF1 loss rescues the reduced OCR caused by loss of COX5A, OPA1, and NDUFAF6. **(A)** Agarose gel electrophoresis of PCR-amplified MTERF1 locus in sg*AAVS1*- or sg*MTERF1*-targeted LHCN-M2 Myoblasts. Clones that matched this targeting were individually validated, and 100% knockout-score clones were pooled for experiments. **(B)** Biological repeats of Fig. 4H-J. (**C)** Mean basal and maximal respiration rates from three biological replicates and three technical replicates in each, obtained from **Fig. S6B and Fig. 4**.

